# Genome-wide data inferring the evolution and population demography of the novel pneumonia coronavirus (SARS-CoV-2)

**DOI:** 10.1101/2020.03.04.976662

**Authors:** Bin Fang, Linlin Liu, Xiao Yu, Xiang Li, Guojun Ye, Juan Xu, Ling Zhang, Faxian Zhan, Guiming Liu, Tao Pan, Yilin Shu, Yongzhong Jiang

**Affiliations:** Influenza Reference Laboratory, Institute of Health Inspection and Testing, Hubei Provincial Center for Disease Control and Prevention, Wuhan 430079, China; Anhui Province Key Laboratory for Conservation and Exploitation of Biological Resource, College of Life Sciences, Anhui Normal University, Wuhu 241000, Anhui, China; Hubei Province Key Laboratory of Occupational Hazard Identification and Control, School of Public Health, Wuhan University of Science and Technology, Wuhan, Hubei, 430065, China; Beijing Agro-Biotechnology Research Center, Beijing Academy of Agriculture and Forestry Sciences, 100097, Beijing, China

**Keywords:** SARS-CoV-2, metagenomic next-generation sequencing, virus evolution, population demography, phylogenetic relationship

## Abstract

As the highly risk and infectious diseases, the outbreak of coronavirus disease 2019 (COVID-19) poses unprecedent challenges to global health. Up to March 3, 2020, SARS-CoV-2 has infected more than 89,000 people in China and other 66 countries across six continents. In this study, we used 10 new sequenced genomes of SARS-CoV-2 and combined 136 genomes from GISAID database to investigate the genetic variation and population demography through different analysis approaches (e.g. Network, EBSP, Mismatch, and neutrality tests). The results showed that 80 haplotypes had 183 substitution sites, including 27 parsimony-informative and 156 singletons. Sliding window analyses of genetic diversity suggested a certain mutations abundance in the genomes of SARS-CoV-2, which may be explaining the existing widespread. Phylogenetic analysis showed that, compared with the coronavirus carried by pangolins (Pangolin-CoV), the virus carried by bats (bat-RaTG13-CoV) has a closer relationship with SARS-CoV-2. The network results showed that SARS-CoV-2 had diverse haplotypes around the world by February 11. Additionally, 16 genomes, collected from Huanan seafood market assigned to 10 haplotypes, indicated a circulating infection within the market in a short term. The EBSP results showed that the first estimated expansion date of SARS-CoV-2 began from 7 December 2019.

## Introduction

As the largest non-segmented genomes among all the RNA viruses (about 30 kb in length), Coronaviruses (CoVs) own the plasticity due to the mutation and recombination, which increased the potential risks of spread across species (Cui and Shi, 2019; Gorbalenya et al., 2020). The COVID-19 (original named 2019-nCOV) is the seventh member of enveloped RNA coronavirus (subgenus, Sarbecovirus; subfamily, Orthocoronavirinae) (Zhu et al., 2020). In early December, 2019, an unexplained pneumonia associated with the severe acute respiratory syndrome coronavirus 2 (SARS-CoV-2) emerged in Wuhan, China (Huang et al., 2020; Lu et al., 2020). The rapid emergence and spread of the SARS-CoV-2 between infected and healthy people became so devastating as a large population within Wuhan was getting infected (Guan et al., 2020; Yang et al., 2020). There’s evidence that SARS-CoV-2 may have originated from bats, but there is no clear information about the intermediate host that transferred it to humans (Wassenaar et al., 2020; Zhou et al., 2020; Wu et al., 2020). On 30 January 2020, the World Health Organization (WHO) declared the outbreak of COVID-19 to be a Public Health Emergency of International Concern. Up to March 3, 2020, SARS-CoV-2 has infected more than 89,000 people in China and other 66 countries across the six continents (source: World Health Organization report, accessed March 3th, 2020).

Despite the worldwide rapid spread, the genomic variation dynamics, evolutionary rate, and virus transmission dynamics of SARS-CoV-2 are not yet well understood. In several recent studies, phylogenetic relationships, variations, evolutionary rates, and propagation dynamics were analyzed using limited genomic data from the SARS-CoV-2 (Lu et al., 2020; Zhou et al., 2020; Wu et al., 2020; Benvenuto et al., 2020; Ceraolo et al., 2020; Chen et al., 2020; Paraskevi et al., 2020; Chan et al., 2020a, 2020b; Li et al., 2020a). As the epidemic progresses, many research institutes around the world have obtained and submitted SARS-CoV-2 genome sequences, increasing the number of SARS-CoV-2 genome sequences in the Data Centers (i.e, GISAID). Given the extremely rapid spread of SARS-CoV-2, an updated analysis with significantly larger sample sizes by incorporating cases throughout the world is urgently needed. This would allow for more accurate identification of SARS-CoV-2 variant dynamics, evolutionary rates, virus transmission dynamics, and epidemic history in order to effectively implement public health measures, promote drugs and vaccines development and to efficiently prevent similar epidemics in the future.

Recent research findings have suggested bat and other non-bat intermediate mammals (such as pangolin) as potential intermediate host for the SARS-CoV-2 that have so far been widely transmitted to humans (Lu et al., 2020; Zhou et al., 2020; Lam et al., 2020). According to the medical information of the first patients to be infected in Wuhan, 27 out of 41 patients were found to be traders of wild mammals in the Huanan seafood market, which directly suggests that the Huanan seafood market as the possible source of origin of the SARS-CoV-2, and later got transmitted it to other areas by the infected people (Huang et al., 2020; Li et al., 2020b). However, due to some infected persons having no relation or access to the Huanan food market, some researchers are skeptical of the market as the only actual origin or the source of SARS-CoV-2 transmission to humans (Cohen, 2020). Therefore, up to date, in the absence of key potential intermediary host, the transmission pattern of SARS-CoV-2 still need to be thoroughly probed for a more reliable and accurate information on the transmission mechanisms of the virus.

In this study, 56 full genome sequences of outgroups and 136 genomes of SARS-CoV-2 from GISAID EpiFluTM database (access date 22 February 2020) were collected. Ten new sequenced genomes of SARS-CoV-2 were obtained by macrogenomic sequencing. The genetic diversity, transmission dynamics and evolutionary history of SARS-CoV-2 were analyzed based on the genomic data to provide understanding on the transmission pathway, and evolutionary characteristics of SARS-CoV-2 outbreak.

## Materials and methods

### Patients and samples

In this study, ten patients with unexplained viral pneumonia from five hospitals in Hubei Province were included. Four samples were collected in December, four in January as well as two samples in February. The December samples were examined by Light Cycle 480II fluorescent PCR instrument (Roche, Basel, Switzerland) for investigating 26 respiratory pathogens and detection of the virus using Severe Acute Respiratory Syndrome Cov (SARS-Cov) and Middle East Respiratory Syndrome Cov (MERS-Cov) primer probes. All the ten samples were detected by the SARS-CoV-2 primer probes. Two samples were used in the virus isolation. The 10 samples were also subjected to metagenomic next-generation sequencing. Specific sample information is presented in Table 1.

**Table 1.** Sample information for 10 SARS-Cov-2 infected patients.

| Sample name | Sample type | Collection date | Collection site | Exposure to huanan seafood marke |
| --- | --- | --- | --- | --- |
| BetaCoV/Wuhan/HBCDC-HB-02/2019 | Bronchoalveolar lavage fluid | 30<sup>th</sup> Dec,2019 | Wuhan, Hubei Province | Yes |
| BetaCoV/Wuhan/HBCDC-HB-01/2019 | Bronchoalveolar lavage fluid | 30<sup>th</sup> Dec,2019 | Wuhan, Hubei Province | Yes |
| BetaCoV/Wuhan/HBCDC-HB-03/2019 | Bronchial scraping | 30<sup>th</sup> Dec,2019 | Wuhan, Hubei Province | Yes |
| BetaCoV/Wuhan/HBCDC-HB-04/2019 | Sputum | 30<sup>th</sup> Dec,2019 | Wuhan, Hubei Province | No |
| BetaCoV/Wuhan/HBCDC-HB-04/2020 | Throat swab | 18<sup>th</sup> Jan,2020 | Wuhan, Hubei Province | No |
| BetaCoV/Wuhan/HBCDC-HB-03/2020 | Throat swab | 18<sup>th</sup> Jan,2020 | Wuhan, Hubei Province | No |
| BetaCoV/Wuhan/HBCDC-HB-02/2020 | Throat swab | 17<sup>th</sup> Jan,2020 | Wuhan, Hubei Province | No |
| BetaCoV/Jingzhou/HBCDC-HB-01/2020 | Throat swab and cultured virus | 8<sup>th</sup> Jan,2020 | Jingzhou,Hubei Province | Yes |
| BetaCoV/Wuhan/HBCDC-HB-06/2020 | Faeces | 7<sup>th</sup> Feb,2020 | Wuhan, Hubei Province | No |
| BetaCoV/Tianmen/HBCDC-HB-07/2020 | Throat swab | 8<sup>th</sup> Feb,2020 | Tianmen, Hubei Province | No |

### Library preparation and sequencing

Total RNA extracted from 10 samples were subjected to metagenomic next generation sequencing testing by EZ1 virus mini kit v.2.0 (955134; Qiage, Heiden, Germany). TruSeq Stranded Total RNA Library Prep Kit (Illumina, San Diego, CA, USA) was used to remove rRNA, reverse transcription and synthesize the double-stranded DNA. The fragmenting, modifying the ends, connecting the joints and enriching of DNA were done by NexteraXT library prepkit (Illumina, San Diego, CA, USA), then the DNA library was obtained. Iseq^TM^ 100 i1 Cartridge, Miseq v2 reagent kit, and High output Reagent Cartridge (illumina, San Diego, CA, USA) were used for deep sequencing in Iseq100, Miseq, and Miniseq platforms (illumina, San Diego, CA, USA), respectively. About 1.5 GB data were obtained for each sample. Using CLC Genomics Workbench 12 and Geneious 12.0.1 software (QIAGEN Bioinformatics, Redwood City, CA, USA) and using reference sequence BetaCoV/Wuhan-Hu-2019 (EPI ISL 402125), the raw sequencing data of Illumina were analyzed.

### Phylogenetic reconstructions

In this study, for probing the evolutionary history of SARS-CoV-2 (Table S1), 136 complete genomes from GISAID (Table S1, access on 22 February, 2020) and our new sequenced 10 genomes of this virus (GISAID, Table S1) were included. In total, 56 outgroups were collected following the previous study (Lu et al., 2020). Mafft v.7.450 was used to align the sequence of SARS-CoV-2 with reference sequences (Katoh et al., 2013). Phylogenetic analyses of the genomes were done with RAxML v.8.2.9 (Stamatakis et al., 2014) in 1000 bootstrap replicates, employing the general time reversible nucleotide substitution model. The tree was visualized with FigTree v.1.4.3 (Rambaut et al., 2016).

### Population structure

On this analysis technique, to precisely decode the evolutionary history of SARS-CoV-2, the genome EPI ISL 402131 (bat-RaTG13-CoV) from GISAID was also included as the outgroup following the previous study (Yu, et al., 2020), because it is the closest sister betacoronavirus to SARS-CoV-2. The 136 complete genome sequences of SARS-CoV-2 and an outgroup (bat-RaTG13-CoV) were aligned using MAFFT then the alignment was manually checked using Geneious. For probing the haplotypes relationships among localities, the Network v.4.6.1 (Fluxus Technology, Suffolk, UK) was used to construct the minimum-spanning network based on the full median-joining algorithm (Bandelt et al., 1999). During the alignment, 10 genomes containing ambiguous sites or “N” bases or more degenerated bases were excluded in this study (Table S1). In addition, 4 genome sequences (EPI ISL 409067, EPI ISL 412981, EPI ISL 407071 and EPI ISL 408489) with one degenerated base were included, which were divided into two sequences without degenerated base. For ensuring the accuracy of this analysis, the 5’UTR and 3’UTR contain missing and ambiguous sites of both regions were excluded in the following alignment analyses. DnaSP v.5.10 (Librado et al., 2009) was used to convert the relevant format. Population genetic indices in was estimated in DnaSP, including nucleotide diversity (π) and haplotype diversity (*Hd*). *F*_ST_ between haplotypes was calculated in Mega 6.0 based on haplotype frequency differences with 10,000 permutations (Tamura et al., 2013). Additionally, a sliding window of 500 bp and steps of 50 bp were used to show nucleotide diversity (π) for the entire alignment this data. Nucleotide diversity for the entire alignment was plotted against midpoint positions of each window. To indicate the relative position of the mutations in the genome, we selected the EPI ISL 402124 sequence as the reference genome.

### Population demography

The alignment was then imported into DnaSP for haplotype analyses. Population size changes were estimated based on a constant population size hypothesis using DnaSP, in combination with neutrality tests (Tajima’s *D* and *Fu*’s Fs). The mismatch distribution was estimated based on the genomes of SARS-CoV-2 in Arlequin to test the hypothesis of recent population growth. The Harpending’s raggedness index (Rag) and the sum of squares deviation (SSD) were used to determine the smoothness of observed mismatch distribution and the degree of fit between observed and simulated data (Harpending, 1994). Because of the sensitivity to demographic changes of neutral tests, Tajima’s *D* (Tajima, 1989) and Fu’s Fs (Fu, 1997) were estimated using 10,000 coalescent simulations to assess the significance in Arlequin. In addition, the Extended Bayesian skyline plot (EBSP) analysis was conducted to examine past population dynamics of SARS-CoV-2 based on 146 genomes by BEAST v.1.8 (Drummond et al., 2012). The divergence times were estimated in BEAST using a Bayesian Markov chain Monte Carlo (MCMC) method with a strict clock. In this study, the substitution rate was set as 0.92×10^−3^ (95% CI, 0.33×10^−3^-1.46×10^−3^) substitution/site/year based on the most recent estimation for SARS-CoV-2 (Yu et al., 2020). The other parameters were set as follows: extended Bayesian skyline process, 10 million MCMC generations, sampling every 1,000^th^ iteration, the initial 25% burn-in. Tracer was used to check the convergence of the MCMC analyses (effective sample size [ESS] values >200).

## Results and discussion

### Genomic variations of SARS-CoV-2

According to the database, the genome size of SARS-CoV-2 varied from 29,409 bp to 29,911 bp. In the network dataset, the aligned matrix was 29,130 bp in length including 183 variable sites, of which 27 were parsimony-informative and 156 were singletons, which were classified as 80 haplotypes (Table S2). Nucleotide diversity (π) was 0.16×10^−3^ ± 0.02×10^−3^ (Standard Deviation, SD) and haplotype diversity (*Hd*) was 0.954 ± 0.013 (SD) and variance of *Hd* was 0.16×10^−3^ (Table S3). In this study, our new sequenced 10 genomes of SARS-CoV-2 represented 10 haplotypes (H3, H8, H10, H14, H16, H28, H29, H30, H76 and H78; Table S1). Among these 10 genomes, four samples were collected from those patients who had been exposed to the Huanan seafood market in Wuhan. Additionally, the HBCDC-HB-03/2019 belonged to the core H3 haplotype and most of them represented new haplotype. Sliding window analysis of SARS-CoV-2 revealed significant regional variation across the alignment (Fig. 1). The plot readily showed a relative high degree of nucleotide variation amongst the aligned SARS-CoV-2 genomes for any given window of 500 bp and steps of 50 bp, with the π ranging from 0.00005 to 0.0014. Additionally, based on the curve, there were several relative high variation area, compared with other fragments (Fig. 1). The *F*_ST_ analyses among 80 haplotypes ranged from 0.00003 (e.g, H3 and H9; H14 and H15; H3 and H20) to 0.00134 (H19 and H80), which indicated genetic differentiation among these haplotypes (Fig. 2, Table S4). Overall, the H19 showed relatively large genetic distance with other haplotypes, while the H3 showed opposite pattern, and was also confirmed by the network figure (Fig. 2, Table S4). Overall, SARS-CoV-2 maintain the relatively rich mutations that has occurred across the world.

**Fig. 1.**
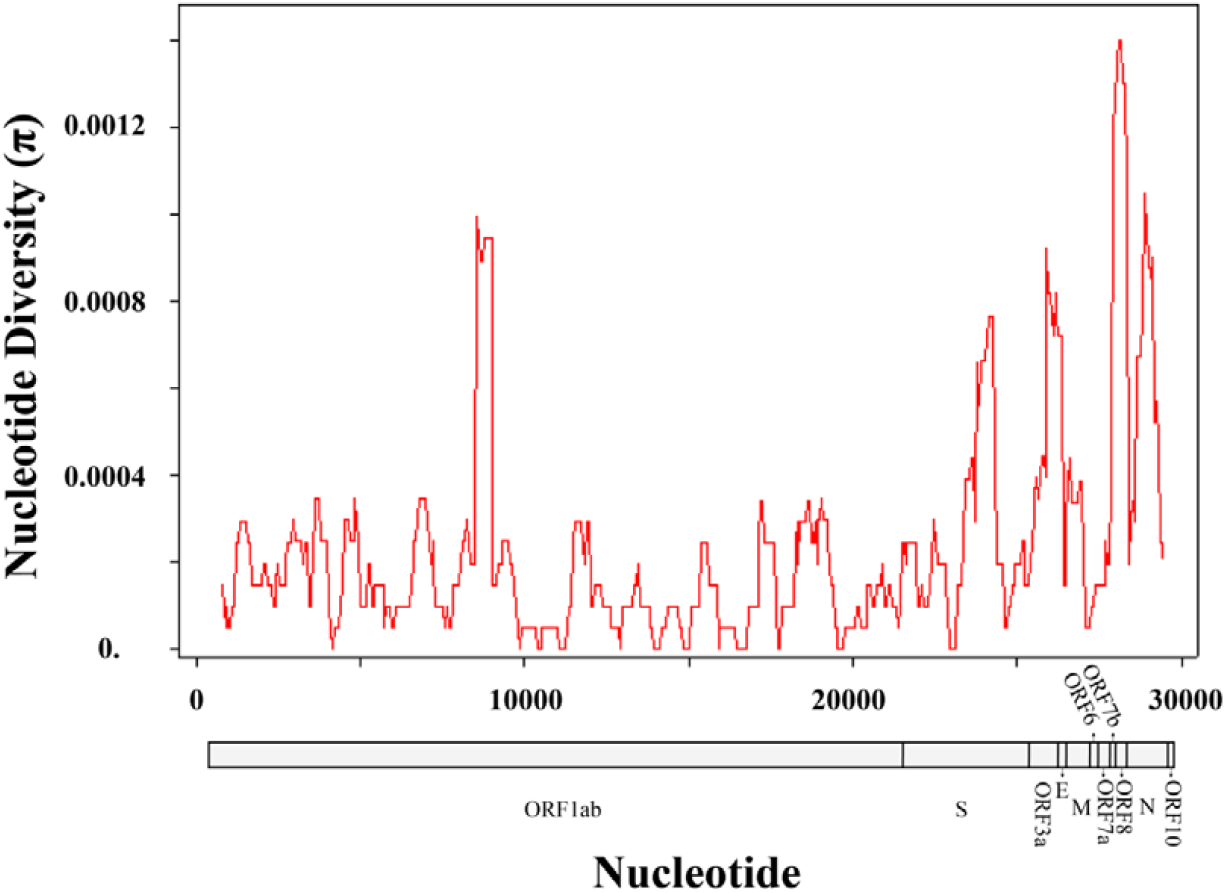
Sliding window analyses showing the nucleotide diversity based on alignment of genomes of SARS-CoV-2. The red line shows the value of nucleotide diversity (π) in a sliding window analysis of window size 500 bp with step size 50, the value is inserted at its mid-point. EPI ISL 402124 sequence was selected as the reference genome to indicate the mutation position.

**Fig. 2.**
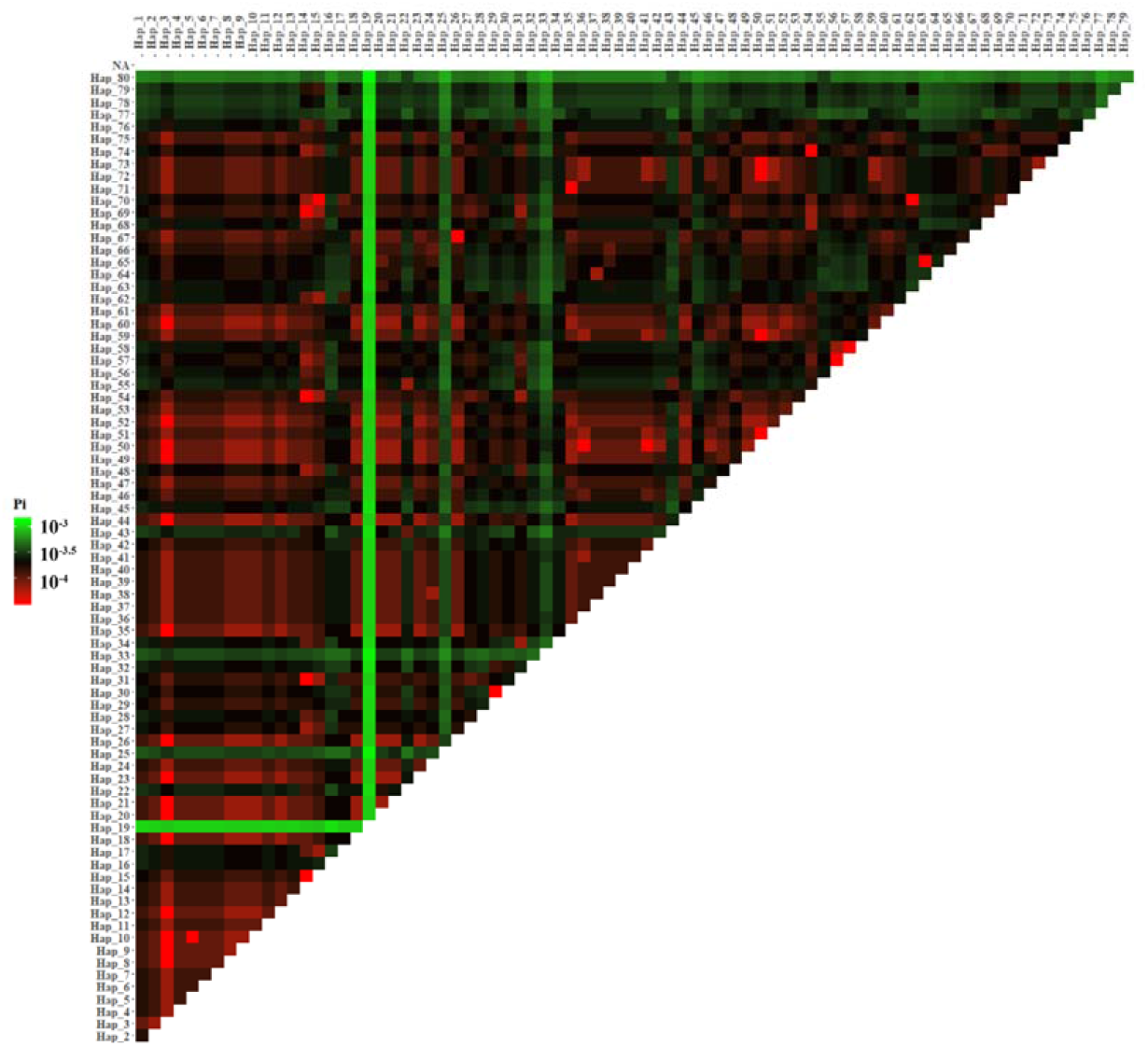
Pairwise *F*_ST_ among 80 haplotypes of SARS-CoV-2. Consistent with the Table S4.

### Phylogenetic relationship of SARS-CoV-2

The virus’s popularity and its intermediate host has been a topic of great concern. Some researchers have suggested that the SARS-CoV-2 originated from the bats (Lu et al., 2020; Zhou et al., 2020). However, some other researchers have proposed that the non-bat intermediate mammals (e.g. pangolins) may be the transmission path of this genus (Lam et al., 2020; Zhang et al., 2020; Xiao et al., 2020). Other studies have also failed to confirm the pangolins as the intermediate host to this viral genus ((Liu et al., 2020). In this study, based on the phylogenetic tree, bat-RaTG13-CoV was the sister group to the SARS-CoV-2 and the two pangolin samples collected from Guangdong which showed a relatively large genetic distance from the SARS-CoV-2 (Fig. 3 and S1). Based on this finding, the study suggested that pangolin may not be the transmission path of the SARS-CoV-2, as had been reiterated by other previous study (Liu et al., 2020). Notably, previous studies have shown that Malayan pangolins could act as potential intermediate host of SARS-CoV-2 (Lam et al., 2020; Xiao et al., 2020). Therefore, more sampling and analysis of the sample across the world may suggest clearer results for more concrete conclusions in future studies.

**Fig. 3.**
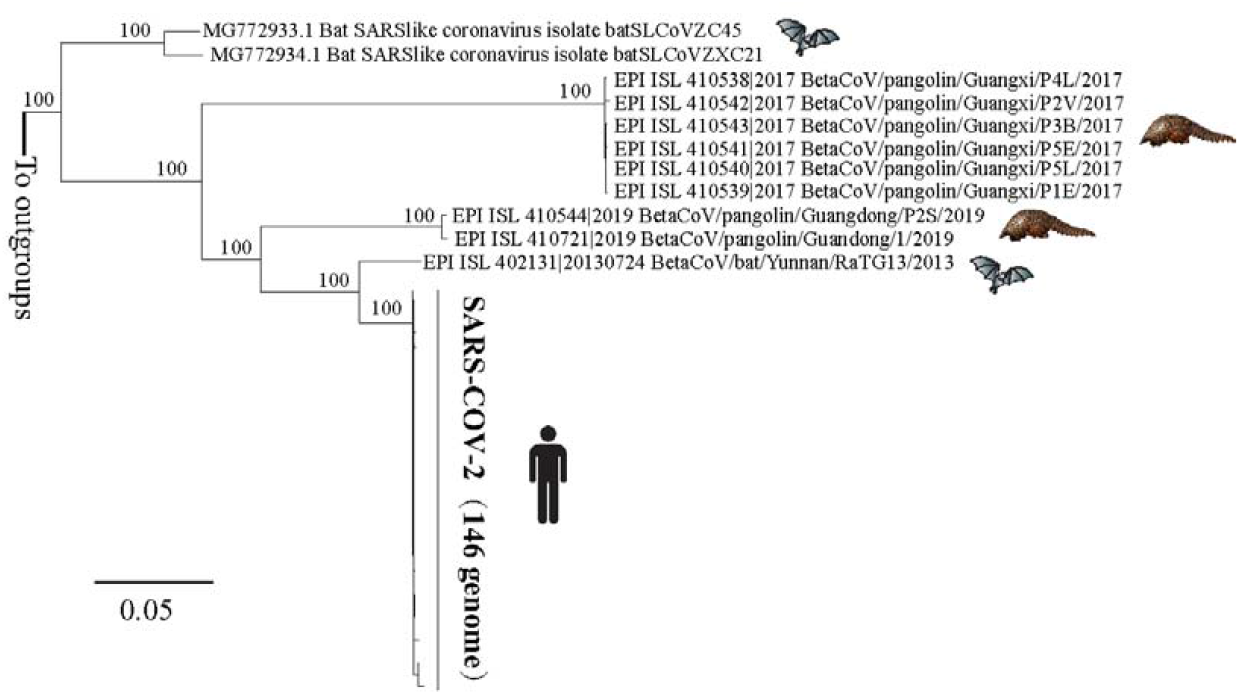
Simplify phylogenetic tree of SARS-CoV-2 form Fig. S1. The nodal numbers are ML bootstrap values.

### Evolutionary relationships of SARS-CoV-2 haplotypes

The evolutionary network of 80 haplotypes of SARS-CoV-2, with bat-RaTG13-CoV as the outgroup, is shown in Figure 4. The network analysis showed typical star network, with several core haplotype node (H3, H14, H15) and edge haplotype nodes. (Fig. 4 and Table S1). In the network, fifty-five satellite haplotypes and H14 connected to the H3 haplotype; eighteen satellite haplotypes and H15+H3 connected to the H14 haplotype; and five satellite haplotypes and H14 connected to H15. Most of these haplogroups were separated by one to six mutations, except two haplotypes (H19 and H80, Fig. 4). The evolutionary network showed that bat-RaTG13-CoV to be connected through a hypothesized haplotype (mv6) to the H15 and H31 haplotypes by single mutations (Fig. 4). However, the mutations between mv6 and bat-RaTG13-CoV was more than 1,200, which indicated that SARS-CoV-2 still has a relative distant kinship with the outgroup (bat-RaTG13-CoV).

**Fig. 4.**
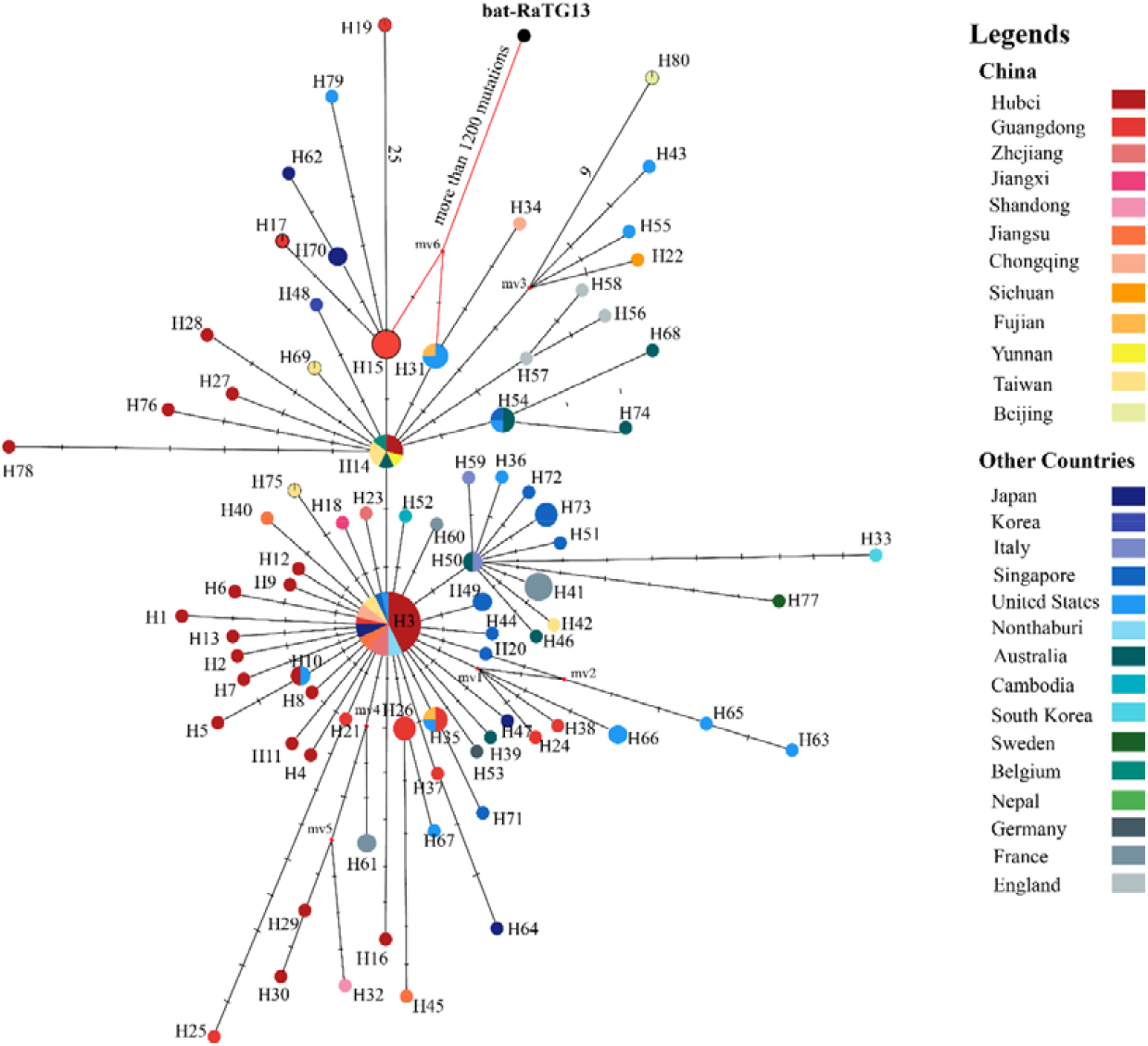
Median-joining network with node sizes proportional to the frequencies of haplotypes in SARS-CoV-2. All the individuals collected before 11^th^ February in 2019. The numbers of mutations separating the haplotypes are shown on the branches, except for the lower than seven-step mutations. The little red diamond nodes indicate undetected haplotypes. The sampling areas are indicated by different colors.

In this study, 16 sequences of Huanan seafood market (4 new sequenced and 12 GISAID data) were collected. Fifteen of the sixteen samples were collected before 1^st^ Januray, 2020 and one samples was collected on 8^th^ Januray, 2020. These samples were represented ten haplotypes (H2, H3, H4, H5, H6, H7, H8, H10, H11, H16), which indicated the high proportion of haplotype diversity in Huanan seafood market. Among those haplotypes, the H3 haplotype, the most abundant, was present in 6 samples from Huanan seafood market, while the other haplotypes were directly derived from the H3 haplotype. All the samples from the Huanan seafood market had the H3 haplotype and other 9 derived haplotypes (Fig. 4), indicating that there were circulated infections within the market in a short term. Additionally, in the network, a total of 65 virus samples from 15 other countries were assigned to 43 haplotypes. Among them, 27 haplotypes were satellite haplotypes of H3 haplotype, 11 haplotypes were satellite haplotypes of H14 haplotype, and 3 haplotypes were satellite haplotype of H15 haplotype. The results showed that SARS-CoV-2 had diverse haplotypes around the world.

Based on the outgroup (bat-RaTG13-CoV) having a possible direct connection with outgroup H15 and H31, it indicates that both the H15 and H31 were the suggested ancestral haplotypes (Fig. 4), as had also been suggested by other previous study (Yu et al., 2020). The H15 was only recovered from five Shenzhen (Guangdong) samples, while H31 included 3 United States samples and 1 Fujian sample from China (Fig. 4). Based on the epidemiological statistics, all the patients from the H15 were ever traveled to Wuhan (Chan et al., 2020b). As to the H31, three genomes from the same patient in Unite State (Holshue et al., 2020) and 1 genome from Fujian of China (Table S1) were included. For the patient in Unite State, he may have infected during the period of visiting his family in China, or was infected in some other place (Yu et al., 2020). Based on the theory in the previous study (Yu et al., 2020), this current study also supported two main evolutionary paths, that the available haplotypes could be from H15 through H14 to H3, or from H31 through H14 to H3 (Fig. 4). Both these two path demonstrate that H14 was the key connection from an ancestral haplotype to H3. However, in H14, only two individuals (EPI ISL 406801 and EPI ISL 412979) were collected from Wuhan and the others were not related to Wuhan. The two individuals (EPI ISL 406801 and EPI ISL 412979) were not directly link with the Huanan seafood market.

Cohen (2020) had argued that Wuhan seafood market may not be source of novel virus spreading globally, or at least not be a single source of SARS-CoV-2 (Cohen et al., 2020). In addition, other studies have shown similar results (Yu et al., 2020; Forster et al., 2020). Due to lacking of early samples and important epidemiological clues across the world, in this study, we only can infer similar conclusions based on the outgroup genome (bat-RaTG13-CoV). On the other hand, some studies had proved that Median-joining network analysis of SARS-CoV-2 genomes is neither phylogenetic nor evolutionary, which indicated that misleads more than illuminates an understanding of the evolutionary history of SARS-CoV-2 in humans (Sánchez-Pacheco et al., 2020). Meantime, sampling bias and incorrect rooting make phylogenetic network may led to the unreliable tracing of SARS-COV-2 infections (Mavian et al., 2020). The outgroup is very distant from current SARS-COV-2 sequences, although approximately 1200 substitutions were observed, there could be more than 1200 mutations actually occurred, thus ancestral inferences using this outgroup could be misleading. Hence, in view of many factors that impact on the accurate origin and evolution of SARS-COV-2, it still requires more reliable data and more powerful analysis methods, indicating the later work direction.

### Population size expansion of SARS-CoV-2

As to population demography of SARS-CoV-2, the EBSP results revealed that it experienced an expansion of effective population size for the last 66 days prior to 11^th^ of February, 2020 (Fig. 5). With the latest one being sampled on 11^th^ February 2020, the first expansion date is estimated to had begun from 7^th^ December 2019. The mismatch in the distribution of the total populations was also clearly shown through unimodal (Fig. 6) and the neutral tests (Tajima’s D and Fu’s Fs) revealed statistically significant population expansion of SARS-CoV-2 (Table S3), which also supported the EBSP results. In addition, with the change of time, multiple evidence (i.e. haplotype number, Tajima’s D, Fu’s FS) indicated a relatively highest population growth at around 21^st^ - 27^th^ of January (Table S3). In early December 2019, the first pneumonia cases of unknown origins (Huang et al., 2020). The previously suggested most recent common ancestor (TMRCA) dates for SARS-CoV-2 at 6th December, 2019) (Li et al., 2020a). Additionally, considering that the virus has longest reported incubation period of up to 24 days (Guan et al., 2020), the virus may had first infected humans in mid to late November, which is basically consistent with the results of previous studies (Yu et al., 2020). Therefore, in this study, EBSP revealed that SARS-CoV-2 experienced an effective population size expansion since 7^th^ December 2019, which was also supported by a star-like network, EBSP, the neutral tests (Fu’s and Tajima’s D test) and mismatch analysis.

**Fig. 5.**
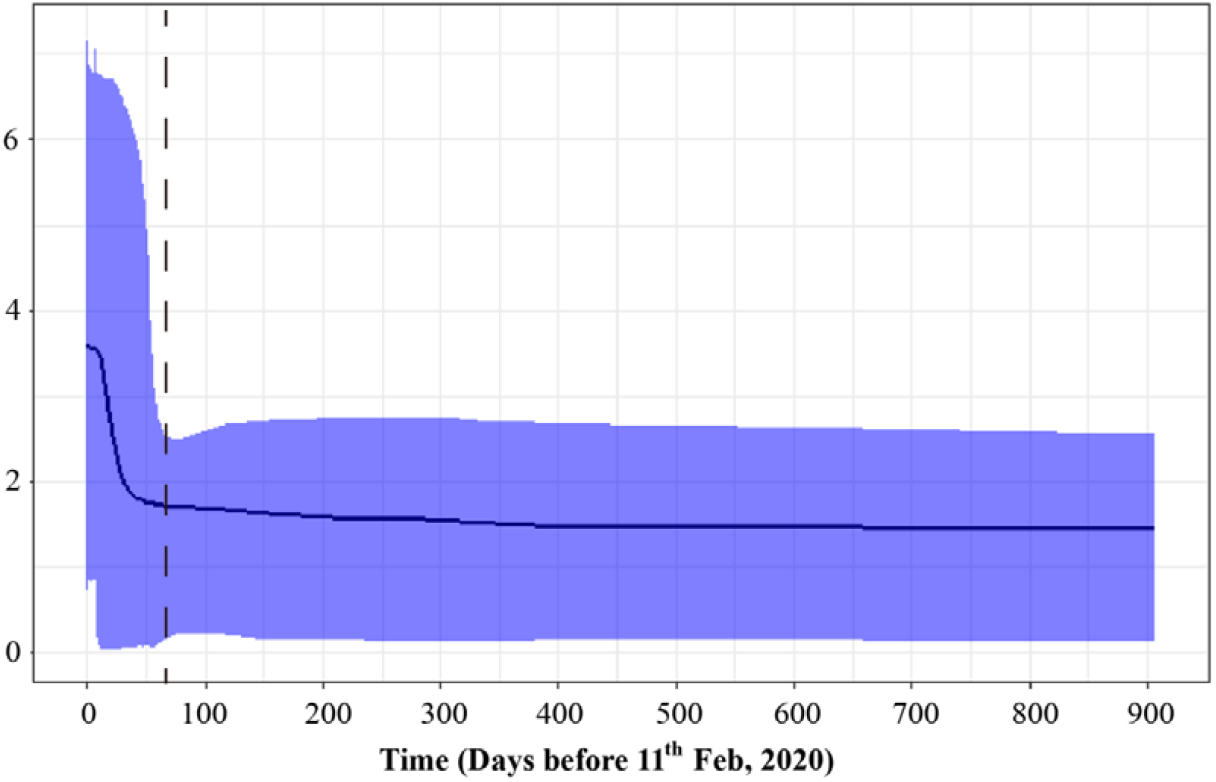
Population fluctuation infered by Extended bayesian skyline plot (EBSP) of SARS-CoV-2. The x-axis indicates time in days BP, and the y-axis indicates the effective population size divided by generation time in units of *N*_eτ_ (the product of effective population size and generation time in days). The blue areas represent 95 % highest posterior density. Time is expressed in days.

**Fig. 6:**
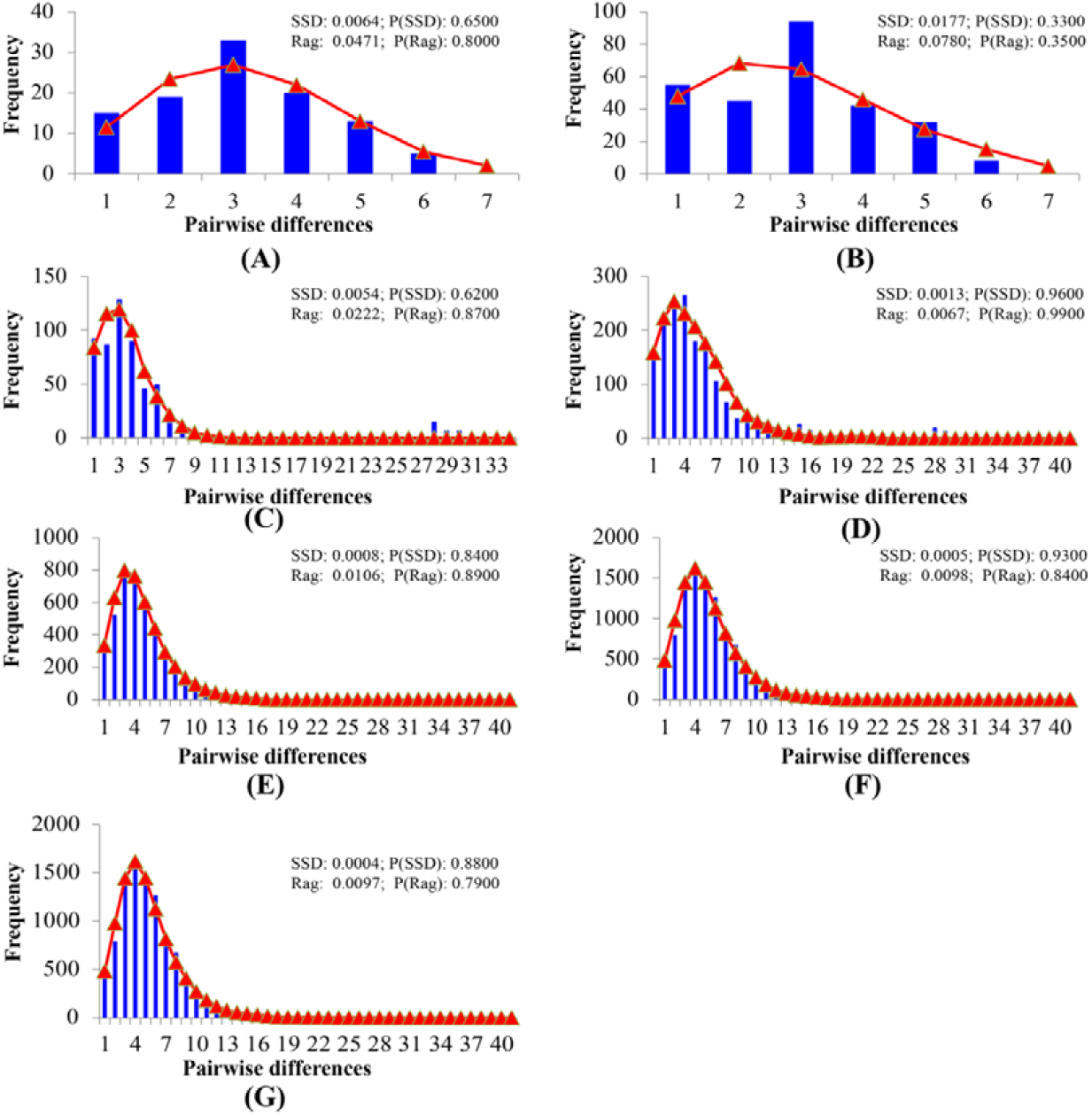
Mismatch distributions analyses for SARS-CoV-2 in different period (A, B, C, D, E, F, G). (A) All the individuals collected during 24^th^ December to 30^th^ December, 2019; (B) 24^th^ December in 2019 to 6^th^ January in 2020; (C) 24^th^ December in 2019 to 13^th^ January in 2020; (D) 24^th^ December in 2019 to 20^th^ January in 2020; (E) 24^th^ December in 2019 to 27^th^ January in 2020; (F) 24^th^ December in 2019 to 3^th^ February in 2020; (G) 24^th^ December in 2019 to 11^th^ February in 2020. The x coordinate represents the number of differences in each pair of sequence comparisons; the y coordinate represents the frequencies of pairwise differences. The blue histogram are the observed frequencies of pairwise divergences among sequences and the red line refers to the expectation under the model of population expansion.

## Conclusion

Coronavirus is a large viruses family, known to cause more serious diseases such as SARS, MERS, etc. SARS-CoV-2 is a new strain of coronavirus which has never been found in human before, which is extremely infectious and harmful to people. In this study, we used 10 new sequenced genomes of SARS-CoV-2 and combined 136 genomes from GISAID database to investigate the genetic variation and population demography. The relative high genetic diversity were disclosed by the mutations abundance, which indicated diverse haplotypes around the world. Compared with the coronavirus carried by pangolins (Pangolin-CoV), the virus carried by bats (bat-RaTG13-CoV) has a closer relationship with SARS-CoV-2. A circulating infection within Huanan seafood market in a short term was disclosed based on multiple haplotypes. Additionally, the first estimated expansion date of SARS-CoV-2 began from 7 December 2019. This study provided data support for understanding the early prevalence of SARS-CoV-2.

## Contributors

Yilin Shu, Tao Pan, and Yongzhong Jiang conceived the research, analyzed the data, interpreted the results, and wrote the draft manuscript; Bin Fang, Linlin Liu, Xiao Yu and Xiang Li performed macrogenome sequencing experiments. Xiao Yu and Xiang Li performed virus isolation experiments. Guojun Ye, Bin Fang, Juan Xu, Ling, Zhang, Yilin Shu and Faxian Zhan collected data. All authors reviewed and approved the final version of the manuscript.

## Acknowledgements

We thank scientists and researchers for depositing the whole genomic sequences of Novel Pneumonia Coronavirus (SARS-CoV-2) in the Global Initiative on Sharing All Influenza Data (GISAID) EpiFlu™; and the GISAID database for allowing us access to sequence for non-commercial scientific research. We also are grateful to Dr. Oscar Omondi Donde (Egerton University, Kenya) for polishing the manuscript language. Thanks to Yifa Zhu, Yangyang Tao and Xierong Li (Tianmen Center for Disease Control and Prevention) for collecting samples and thanks to Maoyi Chen, Jie Hu, Chunlin Mao (Jingzhou Center for Disease Control and Prevention) for sampling.

---

**Fig. S1. Phylogenetic tree of** SARS-CoV-2. The nodal numbers are ML bootstrap values.

