## Supplemental Table 1 for "Genome-wide data inferring the evolution and population demography of the novel pneumonia coronavirus (SARS-CoV-2)"

**Table S1**. Voucher information for 146 genomes of SARS-CoV-2 and bat-RaTG13.

|  | **Accession ID** | **Virus name** | **Location** | **Haplotype** | **Hua Nan Market** | **Collection**  **date** | **Originating lab** | **Submitting lab** | **Authors** |
| --- | --- | --- | --- | --- | --- | --- | --- | --- | --- |
| 1. | EPI_ISL_410984※ | BetaCoV/France/IDF0515-isl/2020 | Europe/France/ Ile-de-France/ Paris |  | Unknown | 2020-01-29 | Department of Infectious and Tropical Diseases, Bichat Claude Bernard Hospital, Paris | National Reference Center for Viruses of Respiratory Infections, Institut Pasteur, Paris | Mélanie Albert, Marion Barbet, Sylvie Behillil, Méline Bizard, Angela Brisebarre, Flora Donati, Vincent Enouf, Maud Vanpeene, Sylvie van der Werf, Yazdan Yazdanpanah, Xavier Lescure |
| 2. | EPI_ISL_410720 | BetaCoV/France/IDF0372-isl/2020 | Europe/France/ Ile-de-France/ Paris | Hap_41 | Unknown | 2020-01-23 | Department of Infectious and Tropical Diseases, Bichat Claude Bernard Hospital, Paris | National Reference Center for Viruses of Respiratory Infections, Institut Pasteur, Paris | Mélanie Albert, Marion Barbet, Sylvie Behillil, Méline Bizard, Angela Brisebarre, Flora Donati, Vincent Enouf, Maud Vanpeene, Sylvie van der Werf, Yazdan Yazdanpanah, Xavier Lescure. |
| 3. | EPI_ISL_410719 | BetaCoV/Singapore/11/2020 | Asia/Singapore | Hap_71 | Unknown | 2020-02-02 | National Public Health Laboratory | National Public Health Laboratory | Octavia S, Mak TM, Cui L, Lin RTP |
| 4. | EPI_ISL_410718 | BetaCoV/Australia/QLD04/2020 | Oceania/Australia/ Queensland/Gold Coast | Hap_54 | Unknown | 2020-02-05 | Pathology Queensland | Public Health Virology Laboratory | Ben Huang, Alyssa Pyke, Amanda De Jong, Andrew Van Den Hurk, Carmel Taylor, David Warrilow, Doris Genge, Elisabeth Gamez, Glen Hewitson, Ian Maxwell Mackay, Inga Sultana, Jamie McMahon, Jean Barcelon, Judy Northill, Mitchell Finger, Natalie Simpson, Neelima Nair, Peter Burtonclay, Peter Moore, Sarah Wheatley, Sean Moody, Sonja Hall-Mendelin, Timothy Gardam, and Frederick Moore. |
| 5. | EPI_ISL_410717 | BetaCoV/Australia/QLD03/2020 | Oceania/Australia/ Queensland/Gold Coast | Hap_74 | Unknown | 2020-02-05 | Pathology Queensland | Public Health Virology Laboratory | Ben Huang, Alyssa Pyke, Amanda De Jong, Andrew Van Den Hurk, Carmel Taylor, David Warrilow, Doris Genge, Elisabeth Gamez, Glen Hewitson, Ian Maxwell Mackay, Inga Sultana, Jamie McMahon, Jean Barcelon, Judy Northill, Mitchell Finger, Natalie Simpson, Neelima Nair, Peter Burtonclay, Peter Moore, Sarah Wheatley, Sean Moody, Sonja Hall-Mendelin, Timothy Gardam, and Frederick Moore. |
| 6. | EPI_ISL_410716 | BetaCoV/Singapore/10/2020 | Asia/Singapore | Hap_73 | Unknown | 2020-02-04 | National Public Health Laboratory, National Centre for Infectious Diseases | National Public Health Laboratory, National Centre for Infectious Diseases | Octavia S, Mak TM, Cui L, Lin RTP |
| 7. | EPI_ISL_410715 | BetaCoV/Singapore/9/2020 | Asia/Singapore | Hap_73 | Unknown | 2020-02-04 | National Public Health Laboratory, National Centre for Infectious Diseases | National Public Health Laboratory, National Centre for Infectious Diseases | Octavia S, Mak TM, Cui L, Lin RTP |
| 8. | EPI_ISL_410714 | BetaCoV/Singapore/8/2020 | Asia/Singapore | Hap_72 | Unknown | 2020-02-03 | National Public Health Laboratory, National Centre for Infectious Diseases | National Public Health Laboratory, National Centre for Infectious Diseases | Octavia S, Mak TM, Cui L, Lin RTP |
| 9. | EPI_ISL_410713 | BetaCoV/Singapore/7/2020 | Asia/Singapore | Hap_51 | Unknown | 2020-01-27 | National Public Health Laboratory, National Centre for Infectious Diseases | National Public Health Laboratory, National Centre for Infectious Diseases | Octavia S, Mak TM, Cui L, Lin RTP |
| 10. | EPI_ISL_410546 | BetaCoV/Italy/INMI1-cs/2020 | Europe/Italy/ Rome | Hap_50 | Unknown | 2020-01-31 | INMI Lazzaro Spallanzani IRCCS | Laboratory of Virology, INMI Lazzaro Spallanzani IRCCS | Maria R. Capobianchi, Cesare E. M. Gruber, Martina Rueca, Fabrizio Carletti, Barbara Bartolini, Francesco Messina, Emanuela Giombini, Francesca Colavita, Concetta Castilletti, Eleonora Lalle, Emanuele Nicastri, Giuseppe Ippolito. |
| 11. | EPI_ISL_410545 | BetaCoV/Italy/INMI1-isl/2020 | Europe/Italy/ Rome | Hap_59 | Unknown | 2020-01-31 | INMI Lazzaro Spallanzani IRCCS | Laboratory of Virology, INMI Lazzaro Spallanzani IRCCS | Maria R. Capobianchi, Cesare E. M. Gruber, Martina Rueca, Fabrizio Carletti, Barbara Bartolini, Francesco Messina, Emanuela Giombini, Francesca Colavita, Concetta Castilletti, Eleonora Lalle, Emanuele Nicastri, Giuseppe Ippolito. |
| 12. | EPI_ISL_410537 | BetaCoV/Singapore/6/2020 | Asia/Singapore | Hap_49 | Unknown | 2020-02-09 | Singapore General Hospital, Molecular Laboratory, Division of Pathology | Programme in Emerging Infectious Diseases, Duke-NUS Medical School | Danielle E Anderson, Martin Linster, Yan Zhuang, Jayanthi Jayakumar, Kian Sing Chan, Lynette LE Oon, Shirin Kalimuddin, Jenny GH Low, Yvonne CF Su, Gavin JD Smith |
| 13. | EPI_ISL_410536 | BetaCoV/Singapore/5/2020 | Asia/Singapore | Hap_73 | Unknown | 2020-02-06 | Singapore General Hospital, Molecular Laboratory, Division of Pathology | Programme in Emerging Infectious Diseases, Duke-NUS Medical School | Danielle E Anderson, Martin Linster, Yan Zhuang, Jayanthi Jayakumar, Kian Sing Chan, Lynette LE Oon, Shirin Kalimuddin, Jenny GH Low, Yvonne CF Su, Gavin JD Smith |
| 14. | EPI_ISL_410535 | BetaCoV/Singapore/4/2020 | Asia/Singapore | Hap_54 | Unknown | 2020-02-03 | National Centre for Infectious Diseases | Programme in Emerging Infectious Diseases, Duke-NUS Medical School | Danielle E Anderson, Martin Linster, Yan Zhuang, Jayanthi Jayakumar, David CB Lye, Yee Sin Leo, Barnaby E Young, Yvonne CF Su, Gavin JD Smith |
| 15. | EPI_ISL_410532 | BetaCoV/Japan/OS-20-07-1/2020 | Asia/Japan/ Osaka | Hap_3 | Unknown | 2020-01-23 | Dept. of Pathology, National Institute of Infectious Diseases | Pathogen Genomics Center, National Institute of Infectious Diseases | Tsuyoshi Sekizuka, Harutaka Katano, Shutoku Matsuyama, Naganori Nao, Kazuya Shirato, Motoi Suzuki, Hideki Hasegawa, Takaji Wakita, Makoto Takeda, Tadaki Suzuki, Makoto Kuroda |
| 16. | EPI_ISL_410531 | BetaCoV/Japan/NA-20-05-1/2020 | Asia/Japan / Nara | Hap_3 | Unknown | 2020-01-25 | Dept. of Pathology, National Institute of Infectious Diseases | Pathogen Genomics Center, National Institute of Infectious Diseases | Tsuyoshi Sekizuka, Harutaka Katano, Shutoku Matsuyama, Naganori Nao, Kazuya Shirato, Motoi Suzuki, Hideki Hasegawa, Takaji Wakita, Makoto Takeda, Tadaki Suzuki, Makoto Kuroda |
| 17. | EPI_ISL_410486  ※ | BetaCoV/France/RA739/2020 | Europe / France / Rhone-Alpes/ Contamines |  | Unknown | 2020-02-25 | CNR Virus des Infections Respiratoires - France SUD | CNR Virus des Infections Respiratoires - France SUD | Bal, Antonin; Destras, Gregory; Gaymard, Alexandre; Bouscambert-Duchamp, Maude; Cheynet, Valérie; Brengel-Pesce, Karen; Morfin-Sherpa, Florence; Valette, Martine; Josset, Laurence; Lina, Bruno. |
| 18. | EPI_ISL_410301 | BetaCoV/Nepal/61/2020 | Asia/Nepal/ Kathmandu | Hap_20 | Unknown | 2020-01-13 | National Influenza Centre, National Public Health Laboratory, Kathmandu, Nepal | The University of Hong Kong | Ranjit Sah , Runa Jha, Daniel Chu, Haogao Gu, Malik Peiris, Anup Bastola, Alfonso J. Rodriguez-Morales, Bibek Kumar Lal, Basu Dev Pandey, Leo Poon |
| 19. | EPI_ISL_410218 | BetaCov/Taiwan/NTU02/2020 | Asia/Taiwan/ Taipei | Hap_75 | Unknown | 2020-02-05 | Department of Laboratory Medicine, National Taiwan University Hospital | Microbial Genomics Core Lab, National Taiwan University Centers of Genomic and Precision Medicine | Shiou-Hwei Yeh, You-Yu Lin, Ya-Yun Lai, Chiao-Ling Li, Shan-Chwen Chang, Pei-Jer Chen, Sui-Yuan Chang |
| 20. | EPI_ISL_410045 | BetaCoV/USA/IL2/2020 | North America/ USA /Illinois | Hap_55 | Unknown | 2020-01-28 | IL Department of Public Health Chicago Laboratory | Pathogen Discovery, Respiratory Viruses Branch, Division of Viral Diseases, Centers for Dieases Control and Prevention | Yan Li, Jing Zhang, Krista Queen, Ying Tao, Anna Uehara, Clinton R. Paden, Xiaoyan Lu, Brian Lynch, Senthil Kumar K. Sakthivel, Brett L. Whitaker, Shifaq Kamili, Lijuan Wang, Janna' R. Murray, Susan I. Gerber, Stephen Lindstrom, Suxiang Tong |
| 21. | EPI_ISL_410044 | BetaCoV/USA/CA6/2020 | North America/ USA /California | Hap_3 | Unknown | 2020-01-27 | California Department of Public Health | Pathogen Discovery, Respiratory Viruses Branch, Division of Viral Diseases, Centers for Dieases Control and Prevention | Jing Zhang, Krista Queen, Yan Li, Ying Tao, Anna Uehara, Clinton R. Paden, Xiaoyan Lu, Brian Lynch, Senthil Kumar K. Sakthivel, Brett L. Whitaker, Shifaq Kamili, Lijuan Wang, Janna' R. Murray, Susan I. Gerber, Stephen Lindstrom, Suxiang Tong |
| 22. | EPI_ISL_409067 | BetaCoV/USA/MA1/2020 | North America/ USA/  Massachusetts | Hap_63 | Unknown | 2020-01-29 | Massachusetts Department of Public Health | Pathogen Discovery, Respiratory Viruses Branch, Division of Viral Diseases, Centers for Dieases Control and Prevention | Clinton R. Paden, Jing Zhang, Krista Queen, Yan Li, Ying Tao, Anna Uehara, Xiaoyan Lu, Brian Lynch, Senthil Kumar K. Sakthivel, Brett L. Whitaker, Shifaq Kamili, Lijuan Wang, Janna' R. Murray, Susan I. Gerber, Stephen Lindstrom, Suxiang Tong |
| 23. | EPI_ISL_408978  ※ | BetaCoV/Wuhan/WH05/2020 | Asia/China/Hubei/ Wuhan |  | Unknown | 2020-02-07 | Wuhan Fourth Hospital | Beijing Genomics Institute (BGI) | Weijun Chen |
| 24. | EPI_ISL_408977 | BetaCoV/Sydney/3/2020 | Oceania/ Australia/New South Wales/ Sydney | Hap_50 | Unknown | 2020-01-25 | Serology, Virology and OTDS Laboratories (SAViD), NSW Health Pathology Randwick | NSW Health Pathology - Institute of Clinical Pathology and Medical Research; Centre for Infectious Diseases and Microbiology Laboratory Services; Westmead Hospital; University of Sydney | Eden J-S, Carter I, Rahman H, Rawlinson W, Holmes EC, Rockett R, O’Sullivan MV, Sintchenko V, Chen SC, Maddocks S, Kok J and Dwyer DE for the 2019-nCoV Study Group |
| 25. | ISL_408976 | BetaCoV/Sydney/2/2020 | Oceania/ Australia/New South Wales/ Sydney | Hap_39 | Unknown | 2020-01-22 | Centre for Infectious Diseases and Microbiology Laboratory Services | NSW Health Pathology - Institute of Clinical Pathology and Medical Research; Westmead Hospital; University of Sydney | Rockett R, Sadsad R, Eden J-S, Carter I, Rahman H, Holmes EC, O’Sullivan MV, Sintchenko V, Chen SC, Maddocks S, Kok J and Dwyer DE for the 2019-nCoV Study Group |
| 26. | EPI_ISL_408670 | BetaCoV/USA/WI1/2020 | North America/ USA / Wisconsin | Hap_35 | Unknown | 2020-01-31 | Wisconsin Department of Health Services | Pathogen Discovery, Respiratory Viruses Branch, Division of Viral Diseases, Centers for Dieases Control and Prevention | Jing Zhang, Anna Uehara, Krista Queen, Yan Li, Ying Tao, Clinton R. Paden, Xiaoyan Lu, Brian Lynch, Senthil Kumar K. Sakthivel, Brett L. Whitaker, Shifaq Kamili, Lijuan Wang, Janna' R. Murray, Susan I. Gerber, Stephen Lindstrom, Suxiang Tong |
| 27. | EPI_ISL_408669 | BetaCoV/Japan/KY-V-029/2020 | Asia/Japan/ Kyoto | Hap_64 | Unknown | 2020-01-29 | Dept. of Virology III, National Institute of Infectious Diseases | Pathogen Genomics Center, National Institute of Infectious Diseases | Tsuyoshi Sekizuka, Shutoku Matsuyama, Naganori Nao, Kazuya Shirato, Makoto Takeda, Makoto Kuroda |
| 28. | EPI_ISL_408667 | BetaCoV/Japan/TY-WK-521/2020 | Asia/Japan/ Tokyo | Hap_70 | Unknown | 2020-01-31 | Dept. of Virology III, National Institute of Infectious Diseases | Pathogen Genomics Center, National Institute of Infectious Diseases | Tsuyoshi Sekizuka, Shutoku Matsuyama, Naganori Nao, Kazuya Shirato, Makoto Takeda, Makoto Kuroda |
| 29. | EPI_ISL_408666 | BetaCoV/Japan/TY-WK-501/2020 | Asia/Japan/ Tokyo | Hap_70 | Unknown | 2020-01-31 | Dept. of Virology III, National Institute of Infectious Diseases | Pathogen Genomics Center, National Institute of Infectious Diseases | Tsuyoshi Sekizuka, Shutoku Matsuyama, Naganori Nao, Kazuya Shirato, Makoto Takeda, Makoto Kuroda |
| 30. | EPI_ISL_408665 | BetaCoV/Japan/TY-WK-012/2020 | Asia/Japan/ Tokyo | Hap_62 | Unknown | 2020-01-29 | Dept. of Virology III, National Institute of Infectious Diseases | Pathogen Genomics Center, National Institute of Infectious Diseases | Tsuyoshi Sekizuka, Shutoku Matsuyama, Naganori Nao, Kazuya Shirato, Makoto Takeda, Makoto Kuroda |
| 31. | EPI_ISL_408515 | BetaCoV/Wuhan/IVDC-HB-envF13-21/2020 | Asia/China/ Hubei / Wuhan | Hap_13 | Unknown | 2020-01-01 | Institute of Viral Disease Control and Prevention, China CDC | Institute of Viral Disease Control and Prevention, China CDC | William J. Liu, Peipei Liu, Xiang Zhao, Peihua Niu, Yingze Zhao, Wenwen Lei, Ziqian Xu, Shumei Zou, Wei Zhen, Beiwei Ye, Mengjie Yang, Weifeng Shi, Roujian Lu, Wenjie Tan, Zhixiao Chen, Yuchao Wu, Juan Song, Weimin Zhou, Dayan Wang, Jun Han, Wenbo Xu, George F. Gao, Guizhen Wu |
| 32. | EPI_ISL_408514 | BetaCoV/Wuhan/IVDC-HB-envF13-20/2020 | Asia/China/ Hubei/Wuhan | Hap_3 | Unknown | 2020-01-01 | Institute of Viral Disease Control and Prevention, China CDC | Institute of Viral Disease Control and Prevention, China CDC | William J. Liu, Peipei Liu, Xiang Zhao, Peihua Niu, Yingze Zhao, Wenwen Lei, Ziqian Xu, Shumei Zou, Wei Zhen, Beiwei Ye, Mengjie Yang, Weifeng Shi, Roujian Lu, Wenjie Tan, Zhixiao Chen, Yuchao Wu, Juan Song, Weimin Zhou, Dayan Wang, Jun Han, Wenbo Xu, George F. Gao, Guizhen Wu |
| 33. | EPI_ISL_408512  ※ | BetaCoV/Wuhan/IVDC-HB-envF54/2020 | Asia/China/ Hubei / Wuhan |  | Unknown | 2020-01-01 | Institute of Viral Disease Control and Prevention, China CDC | Institute of Viral Disease Control and Prevention, China CDC | William J. Liu, Peipei Liu, Xiang Zhao, Peihua Niu, Yingze Zhao, Wenwen Lei, Ziqian Xu, Beiwei Ye, Weifeng Shi, Roujian Lu, Wenjie Tan, Zhixiao Chen, Yuchao Wu, Juan Song, Dayan Wang, Jun Han, Wenbo Xu, George F. Gao, Guizhen Wu |
| 34. | EPI_ISL_408511  ※ | BetaCoV/Wuhan/IVDC-HB-envF13/2020 | Asia/China/ Hubei / Wuhan |  | Unknown | 2020-01-01 | Institute of Viral Disease Control and Prevention, China CDC | Institute of Viral Disease Control and Prevention, China CDC | William J. Liu, Peipei Liu, Xiang Zhao, Peihua Niu, Yingze Zhao, Wenwen Lei, Ziqian Xu, Beiwei Ye, Weifeng Shi, Roujian Lu, Wenjie Tan, Zhixiao Chen, Yuchao Wu, Juan Song, Dayan Wang, Jun Han, Wenbo Xu, George F. Gao, Guizhen Wu |
| 35. | EPI_ISL_408489 | BetaCoV/Taiwan/NTU01/2020 | Asia/Taiwan/ Taipei | Hap_14 | Unknown | 2020-01-31 | Department of Laboratory Medicine, National Taiwan University Hospital | Microbial Genomics Core Lab, National Taiwan University Centers of Genomic and Precision Medicine | Shiou-Hwei Yeh, You-Yu Lin, Ya-Yun Lai, Chiao-Ling Li, Shan-Chwen Chang, Pei-Jer Chen, Sui-Yuan Chang |
| 36. | EPI_ISL_408488 | BetaCoV/Jiangsu/IVDC-JS-001/2020 | Asia/China/ Jiangsu /Huaian | Hap_3 | Unknown | 2020-01-19 | National Institute for Viral Disease Control and Prevention, China CDC | National Institute for Viral Disease Control & Prevention, CCDC | Wenjie Tan, Shenjiao Wang, Wenling Wang, Peihua Niu, Roujian Lu, Kangchen Zhao, Xiang Zhao, Baoying Huang, Li Zhao, Fei Ye, Wenbo Xu, George F. Gao, Guizhen Wu |
| 37. | EPI_ISL_408487  ※ | BetaCoV/Henan/IVDC-HeN-002/2020 | Asia/China/ Henan/ Zhengzhou |  | Unknown | 2020-01-20 | National Institute for Viral Disease Control and Prevention, China CDC | National Institute for Viral Disease Control & Prevention, China CDC | Wenjie Tan, Jin Xu, Wenling Wang, Peihua Niu, Roujian Lu, Xueyong Huang, Xiang Zhao, Baoying Huang, Li Zhao, Fei Ye, Wenbo Xu, George F. Gao, Guizhen Wu |
| 38. | EPI_ISL_408486 | BetaCoV/Jiangxi/IVDC-JX-002/2020 | Asia/China/ Jiangxi/ Pingxiang | Hap_18 | Unknown | 2020-01-11 | National Institute for Viral Disease Control and Prevention, China CDC | National Institute for Viral Disease Control & Prevention, CCDC | Wenjie Tan, Yong Shi, Wenling Wang, Peihua Niu, Roujian Lu, Jianxiong Li, Xiang Zhao, Baoying Huang, Li Zhao, Fei Ye, Wenbo Xu, George F. Gao, Guizhen Wu |
| 39. | EPI_ISL_408485 | BetaCoV/Beijing/IVDC-BJ-005/2020 | Asia/China/ Beijing | Hap_80 | Unknown | 2020-01-18 | National Institute for Viral Disease Control and Prevention, China CDC | National Institute for Viral Disease Control & Prevention, CCDC | Wenjie Tan,Quanyi Wang,Wenling Wang, Peihua Niu,Roujian Lu,Yang Pan,Xiang Zhao,Baoying Huang,Li Zhao,Fei Ye,Wenbo Xu,George F. Gao,Guizhen Wu |
| 40. | EPI_ISL_408484 | BetaCoV/Sichuan/IVDC-SC-001/2020 | Asia/China/ Sichuan/Chengdu | Hap_22 | Unknown | 2020-01-15 | National Institute for Viral Disease Control and Prevention, China CDC | National Institute for Viral Disease Control & Prevention, CCDC | Wenjie Tan, Jianan Xu, Wenling Wang, Peihua Niu, Roujian Lu, Huiping Yang, Xiang Zhao, Baoying Huang, Li Zhao, Fei Ye, Wenbo Xu, George F. Gao, Guizhen Wu |
| 41. | EPI_ISL_408483  ※ | BetaCoV/Shanghai/IVDC-SH-001/2020 | Asia/China/ Shanghai |  | Unknown | 2020-01-20 | National Institute for Viral Disease Control and Prevention, China CDC | National Institute for Viral Disease Control & Prevention, CCDC | Wenjie Tan，Zhen Teng,Xiang Zhao，Wenling Wang， Peihua Niu，Roujian Lu,Chongshan Li,Baoying Huang，Li Zhao，Fei Ye，Wenbo Xu，George F. Gao，Guizhen Wu |
| 42. | EPI_ISL_408482 | BetaCoV/Shandong/IVDC-SD-001/2020 | Asia/China/ Shandong/ Qingdao | Hap_32 | Unknown | 2020-01-19 | National Institute for Viral Disease Control and Prevention, China CDC | National Institute for Viral Disease Control & Prevention, CCDC | Wenjie Tan, Zhaoguo Wang, Xiang Zhao, Wenling Wang, Peihua Niu, Roujian Lu, Ti Liu, Baoying Huang, Li Zhao, Fei Ye, Wenbo Xu, George F. Gao, Guizhen Wu |
| 43. | EPI_ISL_408481 | BetaCoV/Chongqing/IVDC-CQ-001/2020 | Asia/China/ Chongqing | Hap_3 | Unknown | 2020-01-18 | National Institute for Viral Disease Control and Prevention, China CDC | National Institute for Viral Disease Control & Prevention, CCDC | Wenjie Tan, Hengqin Wang, Xiang Zhao, Wenling Wang, Peihua Niu, Roujian Lu, Sheng Ye, Baoying Huang, Li Zhao, Fei Ye, Wenbo Xu, George F. Gao, Guizhen Wu |
| 44. | EPI_ISL_408480 | BetaCoV/Yunnan/IVDC-YN-003/2020 | Asia/China/ Yunnan/ Kunming | Hap_14 | Unknown | 2020-01-17 | National Institute for Viral Disease Control and Prevention, China CDC | National Institute for Viral Disease Control & Prevention, CCDC | Wenjie Tan，Xiaoqing Fu，Xiang Zhao，Wenling Wang， Peihua Niu，Roujian Lu,Yanhong Sun，Baoying Huang，Li Zhao，Fei Ye，Wenbo Xu，George F. Gao，Guizhen Wu |
| 45. | EPI_ISL_408479 | BetaCoV/Chongqing/ZX01/2020 | Asia/China/ Chongqing/ Zhongxian | Hap_3 | Unknown | 2020-01-23 | Zhongxian Center for Disease Control and Prevention | Chongqing Municipal Center for Disease Control and Prevention | Ye Sheng, Tang Yun, Ling Hua, Zhang Hong, Yu zhen,Chen Shuang,Tan ZhangPing, Su Kun, Li Qin, Tang Wenge, Rong Rong |
| 46. | EPI_ISL_408478 | BetaCoV/Chongqing/YC01/2020 | Asia/China/ Chongqinq/ Yongchuan | Hap_34 | Unknown | 2020-01-21 | Yongchuan District Center for Disease Control and Prevention | Chongqing Municipal Center for Disease Control and Prevention | Ye Sheng, Tang Yun, Ling Hua,Yu zhen,Chen Shuang,Tan ZhangPing, Su Kun, Li Qing, Tang Wenge, Rong Rong |
| 47. | EPI_ISL_408431 | BetaCov/France/IDF0626/2020 | Europe/France/ Ile-de-France/ Paris | Hap_60 | Unknown | 2020-01-29 | Sorbonne Université, Inserm et Assistance Publique-Hôpitaux de Paris (Pitié Salpétrière) | National Reference Center for Viruses of Respiratory Infections, Institut Pasteur, Paris | Mélanie Albert, Marion Barbet, Sylvie Behillil, Méline Bizard, Angela Brisebarre, Flora Donati, Vincent Enouf, Maud Vanpeene, Sylvie van der Werf, Sonia Burrel, Anne-Geneviève Marcelin, Vincent Calvez, David Boutolleau, Elise Klément, Valérie Pourcher, Eric Caumes. |
| 48. | EPI_ISL_408430 | BetaCoV/France/IDF0515/2020 | Europe/France/ Ile-de-France/ Paris | Hap_61 | Unknown | 2020-01-29 | Department of Infectious and Tropical Diseases, Bichat Claude Bernard Hospital, Paris | National Reference Center for Viruses of Respiratory Infections, Institut Pasteur, Paris | Mélanie Albert, Marion Barbet, Sylvie Behillil, Méline Bizard, Angela Brisebarre, Flora Donati, Vincent Enouf, Maud Vanpeene, Sylvie van der Werf, Yazdan Yazdanpanah, Xavier Lescure |
| 49. | EPI_ISL_408010 | BetaCoV/USA/CA5/2020 | North America/ USA/California | Hap_67 | Unknown | 2020-01-29 | California Department of Health | Pathogen Discovery, Respiratory Viruses Branch, Division of Viral Diseases, Centers for Dieases Control and Prevention | Ying Tao, Krista Queen, Jing Zhang, Yan Li, Anna Uehara, Clinton Paden, Xiaoyan Lu, Brian Lynch, Senthil Kumar K. Sakthivel, Brett L. Whitaker, Shifaq Kamili, Lijuan Wang, Janna' R. Murray, Susan I. Gerber, Stephen Lindstrom, Suxiang Tong |
| 50. | EPI_ISL_408009 | BetaCoV/USA/CA4/2020 | North America/ USA /California | Hap_66 | Unknown | 2020-01-29 | California Department of Health | Pathogen Discovery, Respiratory Viruses Branch, Division of Viral Diseases, Centers for Dieases Control and Prevention | Krista Queen, Jing Zhang, Yan Li, Ying Tao, Anna Uehara, Clinton Paden, Xiaoyan Lu, Brian Lynch, Senthil Kumar K. Sakthivel, Brett L. Whitaker, Shifaq Kamili, Lijuan Wang, Janna' R. Murray, Susan I. Gerber, Stephen Lindstrom, Suxiang Tong |
| 51. | EPI_ISL_408008 | BetaCoV/USA/CA3/2020 | North America/ USA /California | Hap_66 | Unknown | 2020-01-29 | California Department of Health | Pathogen Discovery, Respiratory Viruses Branch, Division of Viral Diseases, Centers for Disease Control and Prevention | Krista Queen, Jing Zhang, Yan Li, Ying Tao, Anna Uehara, Clinton Paden, Xiaoyan Lu, Brian Lynch, Senthil Kumar K. Sakthivel, Brett L. Whitaker, Shifaq Kamili, Lijuan Wang, Janna' R. Murray, Susan I. Gerber, Stephen Lindstrom, Suxiang Tong |
| 52. | EPI_ISL_407988 | BetaCoV/Singapore/3/2020 | Asia/Singapore | Hap_3 | Unknown | 2020-02-01 | National Centre for Infectious Diseases | Programme in Emerging Infectious Diseases, Duke-NUS Medical School | Danielle E Anderson, Martin Linster, Yan Zhuang, Jayanthi Jayakumar, David CB Lye, Yee Sin Leo, Barnaby E Young, Yvonne CF Su, Linfa Wang, Gavin JD Smith |
| 53. | EPI_ISL_407987 | BetaCoV/Singapore/2/2020 | Asia /Singapore | Hap_49 | Unknown | 2020-02-01 | National Centre for Infectious Diseases | Programme in Emerging Infectious Diseases, Duke-NUS Medical School | Danielle E Anderson, Martin Linster, Yan Zhuang, Jayanthi Jayakumar, David CB Lye, Yee Sin Leo, Barnaby E Young, Yvonne CF Su, Linfa Wang, Gavin JD Smith |
| 54. | EPI_ISL_407976 | BetaCoV/Belgium/GHB-03021/2020 | Europe/Belgium/ Leuven | Hap_14 | Unknown | 2020-02-03 | KU Leuven, Clinical and Epidemiological Virology | KU Leuven, Clinical and Epidemiological Virology | Bert Vanmechelen, Elke Wollants, Annabel Rector, Els Keyaerts, Lies Laenen, Marc Van Ranst, and Piet Maes |
| 55. | EPI_ISL_407896 | BetaCoV/Australia/QLD02/2020 | Oceania/Australia/ Queensland/Gold Coast | Hap_68 | Unknown | 2020-01-30 | Pathology Queensland | Public Health Virology Laboratory | Ben Huang, Alyssa Pyke, Amanda De Jong, Andrew Van Den Hurk, Carmel Taylor, David Warrilow, Doris Genge, Elisabeth Gamez, Glen Hewitson, Ian Maxwell Mackay, Inga Sultana, Jamie McMahon, Jean Barcelon, Judy Northill, Mitchell Finger, Natalie Simpson, Neelima Nair, Peter Burtonclay, Peter Moore, Sarah Wheatley, Sean Moody, Sonja Hall-Mendelin, Timothy Gardam, and Frederick Moore. |
| 56. | EPI_ISL_407894 | BetaCoV/Australia/QLD01/2020 | Oceania/Australia/ Queensland/ Gold Coast | Hap_54 | Unknown | 2020-01-28 | Pathology Queensland | Public Health Virology Laboratory | Ben Huang, Alyssa Pyke, Amanda De Jong, Andrew Van Den Hurk, Carmel Taylor, David Warrilow, Doris Genge, Elisabeth Gamez, Glen Hewitson, Ian Maxwell Mackay, Inga Sultana, Jamie McMahon, Jean Barcelon, Judy Northill, Mitchell Finger, Natalie Simpson, Neelima Nair, Peter Burtonclay, Peter Moore, Sarah Wheatley, Sean Moody, Sonja Hall-Mendelin, Timothy Gardam, and Frederick Moore. |
| 57. | EPI_ISL_407893 | BetaCoV/Australia/NSW01/2020 | Oceania/Australia/New South Wales/ Sydney | Hap_14 | Unknown | 2020-01-24 | Centre for Infectious Diseases and Microbiology Laboratory Services | NSW Health Pathology - Institute of Clinical Pathology and Medical Research; Westmead Hospital; University of Sydney | Eden J-S, Carter I, Rahman H, Holmes EC, Rockett R, O’Sullivan MV, Sintchenko V, Chen SC, Maddocks S, Kok J and Dwyer DE for the 2019-nCoV Study Group |
| 58. | EPI_ISL_407313 | BetaCoV/Hangzhou/HZCDC0001/2020 | Asia/China/ Zhejiang/ Hangzhou | Hap_3 | Unknown | 2020-01-19 | Hangzhou Center for Disease Control and Prevention | Hangzhou Center for Disease Control and Prevention | Jun Li, Haoqiu Wang, Hua Yu, Lingfeng Mao, Xinfen Yu, Zhou Sun, Qingxin Kong, Xin Qian, Shuchang Chen, Xuchu Wang |
| 59. | EPI_ISL_407215 | BetaCoV/USA/WA1-F6/2020 | North America/ USA/Washington | Hap_31 | Unknown | 2020-01-25 | Washington State Department of Health | Pathogen Discovery, Respiratory Viruses Branch, Division of Viral Diseases, Centers for Dieases Control and Prevention | Krista Queen, Azaibi Tamin, Jennifer Harcourt, Ying Tao, Clinton R. Paden, Jing Zhang, Yan Li, Anna Uehara, Xiaoyan Lu, Shifaq Kamili, Rashi Gautam, Haibin Wang, Janna' R. Murray, Susan I. Gerber, Stephen Lindstrom, Natalie Thornburg, Suxiang Tong |
| 60. | EPI_ISL_407214 | BetaCoV/USA/WA1-A12/2020 | North America/ USA/Washington | Hap_31 | Unknown | 2020-01-25 | WA State Department of Health | Pathogen Discovery, Respiratory Viruses Branch, Division of Viral Diseases, Centers for Dieases Control and Prevention | Krista Queen, Azaibi Tamin, Jennifer Harcourt, Ying Tao, Clinton R. Paden, Jing Zhang, Yan Li, Anna Uehara, Xiaoyan Lu, Shifaq Kamili, Rashi Gautam, Haibin Wang, Janna' R. Murray, Susan I. Gerber, Stephen Lindstrom, Natalie Thornburg, Suxiang Tong |
| 61. | EPI_ISL_407193 | BetaCoV/South Korea/KCDC03/2020 | Asia/South Korea/ Gyeonggido | Hap_48 | Unknown | 2020-01-25 | Korea Centers for Disease Control & Prevention (KCDC) Center for Laboratory Control of Infectious Diseases Division of Viral Diseases | Korea Centers for Disease Control & Prevention (KCDC) Center for Laboratory Control of Infectious Diseases Division of Viral Diseases | Jeong-Min Kim, Yoon-Seok Chung, Namjoo Lee, Mi-Seon Kim, SangHee Woo, Hye-Joon Jo, Sehee Park, Heui Man Kim, Myung Guk Han |
| 62. | EPI_ISL_407084 | BetaCoV/Japan/AI/I-004/2020 | Asia/Japan/Aichi | Hap_47 | Unknown | 2020-01-25 | Department of Virology III, National Institute of Infectious Diseases | Pathogen Genomics Center, National Institute of Infectious Diseases | Tsuyoshi Sekizuka, Shutoku Matsuyama, Naganori Nao, Kazuya Shirato, Shinji Watanabe, Makoto Takeda, Makoto Kuroda |
| 63. | EPI_ISL_407079  ※ | BetaCoV/Finland/1/2020 | Europe/Finland/ Lapland |  | Unknown | 2020-01-29 | Lapland Central Hospital | Department of Virology, University of Helsinki and Helsinki University Hospital, Helsinki, Finland | Teemu Smura, Suvi Kuivanen, Hannimari Kallio-Kokko, Olli Vapalahti |
| 64. | EPI_ISL_407073 | BetaCoV/England/02/2020 | Europe/England | Hap_58 | Unknown | 2020-01-29 | Respiratory Virus Unit, Microbiology Services Colindale, Public Health England | Respiratory Virus Unit, Microbiology Services Colindale, Public Health England | Monica Galiano, Shahjahan Miah, Richard Myers, Angie Lackenby, Omolola Akinbami, Tiina Talts, Leena Bhaw, Kirstin Edwards, Jonathan Hubb, Joanna Ellis, Maria Zambon. |
| 65. | EPI_ISL_407071 | BetaCoV/England/01/2020 | Europe/England | Hap_56 | Unknown | 2020-01-29 | Respiratory Virus Unit, Microbiology Services Colindale, Public Health England | Respiratory Virus Unit, Microbiology Services Colindale, Public Health England | Monica Galiano, Shahjahan Miah, Richard Myers, Angie Lackenby, Omolola Akinbami, Tiina Talts, Leena Bhaw, Kirstin Edwards, Jonathan Hubb, Joanna Ellis, Maria Zambon |
| 66. | EPI_ISL_406973 | BetaCoV/Singapore/1/2020 | Asia/Singapore | Hap_44 | Unknown | 2020-01-23 | Singapore General Hospital | National Public Health Laboratory | Mak, TM; Octavia S; Chavatte JM; Zhou, ZY; Cui, L; Lin, RTP |
| 67. | EPI_ISL_406970 | BetaCoV/Hangzhou/HZ-1/2020 | Asia/China/ Zhejiang/ Hangzhou | Hap_3 | Unknown | 2020-01-20 | Hangzhou Center for Disease and Control Microbiology Lab | Hangzhou Center for Disease and Control Microbiology Lab | Yu Hua, Wang Haoqiu, Li Jun, Yu Xinfeng |
| 68. | EPI_ISL_406862 | BetaCoV/Germany/BavPat1/2020 | Europe/Germany/ Bavaria /Munich | Hap_53 | Unknown | 2020-01-28 | Charité Universitätsmedizin Berlin, Institute of Virology; Institut für Mikrobiologie der Bundeswehr, Munich | Charité Universitätsmedizin Berlin, Institute of Virology | Victor M Corman, Julia Schneider, Talitha Veith, Barbara Mühlemann, Markus Antwerpen, Christian Drosten, Roman Wölfel |
| 69. | EPI_ISL_406844 | BetaCoV/Australia/VIC01/2020 | Oceania/Australia/ Victoria/Clayton | Hap_46 | Unknown | 2020-01-25 | Monash Medical Centre | Collaboration between the University of Melbourne at The Peter Doherty Institute for Infection and Immunity, and the Victorian Infectious Disease Reference Laboratory | Caly,L., Seemann,T., Schultz,M., Druce,J. and Taiaroa,G |
| 70. | EPI_ISL_406801 | BetaCov/Wuhan/WH04/2020 | Asia/China/Hubei/ Wuhan | Hap_14 | No | 2020-01-05 | General Hospital of Central Theater Command of People's Liberation Army of China | BGI & Institute of Microbiology, Chinese Academy of Sciences & Shandong First Medical University & Shandong Academy of Medical Sciences & General Hospital of Central Theater Command of People's Liberation Army of China | Weijun Chen, Yuhai Bi, Weifeng Shi and Zhenhong Hu |
| 71. | EPI_ISL_406800 | BetaCov/Wuhan/WH03/2020 | Asia/China/ Hubei / Wuhan | Hap_3 | Yes | 2020-01-01 | General Hospital of Central Theater Command of People's Liberation Army of China | BGI & Institute of Microbiology, Chinese Academy of Sciences & Shandong First Medical University & Shandong Academy of Medical Sciences & General Hospital of Central Theater Command of People's Liberation Army of China | Weijun Chen, Yuhai Bi, Weifeng Shi and Zhenhong Hu |
| 72. | EPI_ISL_406799※ | BetaCov/Wuhan/WH02/2019 | Asia/China/ Wuhan |  | Unknown | 2019-12-31 | General Hospital of Central Theater Command of People's Liberation Army of China | BGI & Institute of Microbiology, Chinese Academy of Sciences & Shandong First Medical University & Shandong Academy of Medical Sciences & General Hospital of Central Theater Command of People's Liberation Army of China | Weijun Chen, Yuhai Bi, Weifeng Shi and Zhenhong Hu |
| 73. | EPI_ISL_406798 | BetaCov/Wuhan/WH01/2019 | Asia/China/Hubei/ Wuhan | Hap_2 | Yes | 2019-12-26 | General Hospital of Central Theater Command of People's Liberation Army of China | BGI & Institute of Microbiology, Chinese Academy of Sciences & Shandong First Medical University & Shandong Academy of Medical Sciences & General Hospital of Central Theater Command of People's Liberation Army of China | Weijun Chen, Yuhai Bi, Weifeng Shi and Zhenhong Hu |
| 74. | EPI_ISL_406717 | BetaCoV/China/WHU02/2020 | Asia/China/ Hubei / Wuhan | Hap_3 | Unknown | 2020-01-02 | unknown | State Key Laboratory of Virology, Wuhan University | Chen,L., Liu,W., Zhang,Q., Xu,K., Ye,G., Wu,W., Sun,Z., Liu,F., Wu,K., Mei,Y., Zhang,W., Chen,Y., Li,Y., Shi,M., Lan,K. and Liu,Y. |
| 75. | EPI_ISL_406716 | BetaCoV/China/WHU01/2020 | Asia/China/ Hubei / Wuhan | Hap_3 | Unknown | 2020-01-02 | unknown | State Key Laboratory of Virology, Wuhan University | Chen,L., Liu,W., Zhang,Q., Xu,K., Ye,G., Wu,W., Sun,Z., Liu,F., Wu,K., Mei,Y., Zhang,W., Chen,Y., Li,Y., Shi,M., Lan,K. and Liu,Y. |
| 76. | EPI_ISL_406597 | BetaCoV/France/IDF0373/2020 | Europe/France/ Ile-de-France/ Paris | Hap_41 | Unknown | 2020-01-23 | Department of Infectious and Tropical Diseases, Bichat Claude Bernard Hospital, Paris | National Reference Center for Viruses of Respiratory Infections, Institut Pasteur, Paris | Mélanie Albert, Marion Barbet, Sylvie Behillil, Méline Bizard, Angela Brisebarre, Flora Donati, Vincent Enouf, Maud Vanpeene, Sylvie van der Werf, Yazdan Yazdanpanah, Xavier Lescure. |
| 77. | EPI_ISL_406596 | BetaCoV/France/IDF0372/2020 | Europe/France/ Ile-de-France/ Paris | Hap_41 | Unknown | 2020-01-23 | Department of Infectious and Tropical Diseases, Bichat Claude Bernard Hospital, Paris | National Reference Center for Viruses of Respiratory Infections, Institut Pasteur, Paris | Mélanie Albert, Marion Barbet, Sylvie Behillil, Méline Bizard, Angela Brisebarre, Flora Donati, Vincent Enouf, Maud Vanpeene, Sylvie van der Werf, Yazdan Yazdanpanah, Xavier Lescure. |
| 78. | EPI_ISL_406595 | BetaCoV/Shenzhen/SZTH-004/2020 | Asia/China/  Guandong/ Shenzhen | Hap_25 | Unknown | 2020-01-16 | Shenzhen Key Laboratory of Pathogen and Immunity, National Clinical Research Center for Infectious Disease, Shenzhen Third People's Hospital | Shenzhen Key Laboratory of Pathogen and Immunity, National Clinical Research Center for Infectious Disease, Shenzhen Third People's Hospital | Yang Yang, Chenguang Shen, Li Xing, Zhixiang Xu, Haixia Zheng, Yingxia Liu |
| 79. | EPI_ISL_406594 | BetaCoV/Shenzhen/SZTH-003/2020 | Asia/China/ Guandong/ Shenzhen | Hap_24 | Unknown | 2020-01-16 | Shenzhen Key Laboratory of Pathogen and Immunity, National Clinical Research Center for Infectious Disease, Shenzhen Third People's Hospital | Shenzhen Key Laboratory of Pathogen and Immunity, National Clinical Research Center for Infectious Disease, Shenzhen Third People's Hospital | Yang Yang, Chenguang Shen, Li Xing, Zhixiang Xu, Haixia Zheng, Yingxia Liu |
| 80. | EPI_ISL_406593 | BetaCoV/Shenzhen/SZTH-002/2020 | Asia/China/ Guandong/ Shenzhen | Hap_15 | Unknown | 2020-01-13 | Shenzhen Key Laboratory of Pathogen and Immunity, National Clinical Research Center for Infectious Disease, Shenzhen Third People's Hospital | Shenzhen Key Laboratory of Pathogen and Immunity, National Clinical Research Center for Infectious Disease, Shenzhen Third People's Hospital | Yang Yang, Chenguang Shen, Li Xing, Zhixiang Xu, Haixia Zheng, Yingxia Liu |
| 81. | EPI_ISL_406592 | BetaCoV/Shenzhen/SZTH-001/2020 | Asia/China/ Guandong/  Shenzhen | Hap_19 | Unknown | 2020-01-13 | Shenzhen Third People's Hospital | Shenzhen Key Laboratory of Pathogen and Immunity, National Clinical Research Center for Infectious Disease,Shenzhen Third People's Hospital | Yang Yang, Chenguang Shen, Li Xing, Zhixiang Xu, Haixia Zheng, Yingxia Liu |
| 82. | EPI_ISL_406538 | BetaCoV/Guangdong/20SF201/2020 | Asia/China/ Guangdong | Hap_3 | Unknown | 2020-01-23 | Guangdong Provincial Center for Diseases Control and Prevention;Guangdong Provincial Institute of Public Health | Guangdong Provincial Center for Diseases Control and Prevention | Min Kang, Jie Wu, Jing Lu, Tao Liu, Baisheng Li, Shujiang Mei, Feng Ruan, Lifeng Lin, Changwen Ke, Haojie Zhong, Yingtao Zhang, Lirong Zou, Xuguang Chen, Qi Zhu, Jianpeng Xiao, Jianxiang Geng, Zhe Liu, Jianxiong Hu, Weilin Zeng, Xing Li, Yuhuang Liao, Xiujuan Tang, Songjian Xiao, Ying Wang, Yingchao Song, Xue Zhuang, Lijun Liang, Guanhao He, Huihong Deng, Tie Song, Jianfeng He, Wenjun Ma |
| 83. | EPI_ISL_406536 | BetaCoV/Foshan/20SF211/2020 | Asia/China/ Guangdong/ Foshan | Hap_35 | Unknown | 2020-01-22 | Guangdong Provincial Center for Diseases Control and Prevention; Guangdong Provincial Public Health | Guangdong Provincial Center for Diseases Control and Prevention | Min Kang, Jie Wu, Jing Lu, Tao Liu, Baisheng Li, Shujiang Mei, Feng Ruan, Lifeng Lin, Changwen Ke, Haojie Zhong, Yingtao Zhang, Lirong Zou, Xuguang Chen, Qi Zhu, Jianpeng Xiao, Jianxiang Geng, Zhe Liu, Jianxiong Hu, Weilin Zeng, Xing Li, Yuhuang Liao, Xiujuan Tang, Songjian Xiao, Ying Wang, Yingchao Song, Xue Zhuang, Lijun Liang, Guanhao He, Huihong Deng, |
| 84. | EPI_ISL_406535 | BetaCoV/Foshan/20SF210/2020 | Asia/China/ Guangdong/ Foshan | Hap_35 | Unknown | 2020-01-22 | Guangdong Provincial Center for Diseases Control and Prevention; Guangdong Provincial Public Health | Guangdong Provincial Center for Diseases Control and Prevention | Min Kang, Jie Wu, Jing Lu, Tao Liu, Baisheng Li, Shujiang Mei, Feng Ruan, Lifeng Lin, Changwen Ke, Haojie Zhong, Yingtao Zhang, Lirong Zou, Xuguang Chen, Qi Zhu, Jianpeng Xiao, Jianxiang Geng, Zhe Liu, Jianxiong Hu, Weilin Zeng, Xing Li, Yuhuang Liao, Xiujuan Tang, Songjian Xiao, Ying Wang, Yingchao Song, Xue Zhuang, Lijun Liang, Guanhao He, Huihong Deng, Tie Song, Jianfeng He, Wenjun Ma |
| 85. | EPI_ISL_406534 | BetaCoV/Foshan/20SF207/2020 | Asia/China/ Guangdong/ Foshan | Hap_38 | Unknown | 2020-01-22 | Guangdong Provincial Center for Diseases Control and Prevention; Guangdong Provincial Public Health | Guangdong Provincial Center for Diseases Control and Prevention | Min Kang, Jie Wu, Jing Lu, Tao Liu, Baisheng Li, Shujiang Mei, Feng Ruan, Lifeng Lin, Changwen Ke, Haojie Zhong, Yingtao Zhang, Lirong Zou, Xuguang Chen, Qi Zhu, Jianpeng Xiao, Jianxiang Geng, Zhe Liu, Jianxiong Hu, Weilin Zeng, Xing Li, Yuhuang Liao, Xiujuan Tang, Songjian Xiao, Ying Wang, Yingchao Song, Xue Zhuang, Lijun Liang, Guanhao He, Huihong Deng, Tie Song, Jianfeng He, Wenjun Ma |
| 86. | EPI_ISL_406533 | BetaCoV/Guangzhou/20SF206/2020 | Asia/China/ Guangdong/ Guangzhou | Hap_37 | Unknown | 2020-01-22 | Guangdong Provincial Center for Diseases Contorl and Prevention; Guangdong Provinical Public Health | Guangdong Provincial Center for Diseases Control and Prevention | Min Kang, Jie Wu, Jing Lu, Tao Liu, Baisheng Li, Shujiang Mei, Feng Ruan, Lifeng Lin, Changwen Ke, Haojie Zhong, Yingtao Zhang, Lirong Zou, Xuguang Chen, Qi Zhu, Jianpeng Xiao, Jianxiang Geng, Zhe Liu, Jianxiong Hu, Weilin Zeng, Xing Li, Yuhuang Liao, Xiujuan Tang, Songjian Xiao, Ying Wang, Yingchao Song, Xue Zhuang, Lijun Liang, Guanhao He, Huihong Deng, Tie Song, Jianfeng He, Wenjun Ma |
| 87. | EPI_ISL_406531 | BetaCoV/Guangdong/20SF174/2020 | Asia/China/ Guangdong/ Zhuhai | Hap_26 | Unknown | 2020-01-22 | Guangdong Provincial Center for Diseases Contorl and Prevention; Guangdong Provinical Public Health | Guangdong Provincial Center for Disease Control and Prevention | Min Kang, Jie Wu, Jing Lu, Tao Liu, Baisheng Li, Shujiang Mei, Feng Ruan, Lifeng Lin, Changwen Ke, Haojie Zhong, Yingtao Zhang, Lirong Zou, Xuguang Chen, Qi Zhu, Jianpeng Xiao, Jianxiang Geng, Zhe Liu, Jianxiong Hu, Weilin Zeng, Xing Li, Yuhuang Liao, Xiujuan Tang, Songjian Xiao, Ying Wang, Yingchao Song, Xue Zhuang, Lijun Liang, Guanhao He, Huihong Deng, Tie Song, Jianfeng He, Wenjun Ma |
| 88. | EPI_ISL_406036 | BetaCoV/USA/CA2/2020 | North America/ USA/California/ Orange County | Hap_36 | Unknown | 2020-01-22 | California Department of Public Health | Pathogen Discovery, Respiratory Viruses Branch, Division of Viral Diseases, Centers for Dieases Control and Prevention | Anna Uehara, Krista Queen, Ying Tao, Yan Li, Clinton R. Paden, Jing Zhang, Xiaoyan Lu, Brian Lynch, Senthil Kumar K. Sakthivel, Brett L. Whitaker, Shifaq Kamili, Lijuan Wang, Janna' R. Murray, Susan I. Gerber, Stephen Lindstrom, Suxiang Tong |
| 89. | EPI_ISL_406034 | BetaCoV/USA/CA1/2020 | North America/ USA/California/ Los Angeles | Hap_43 | Unknown | 2020-01-23 | California Department of Public Health | Pathogen Discovery, Respiratory Viruses Branch, Division of Viral Diseases, Centers for Dieases Control and Prevention | Anna Uehara, Krista Queen, Ying Tao, Yan Li, Clinton R. Paden, Jing Zhang, Xiaoyan Lu, Brian Lynch, Senthil Kumar K. Sakthivel, Brett L. Whitaker, Shifaq Kamili, Lijuan Wang, Janna' R. Murray, Susan I. Gerber, Stephen Lindstrom, Suxiang Tong |
| 90. | EPI_ISL_406031 | BetaCoV/Taiwan/2/2020 | Asia/Taiwan/ Kaohsiung | Hap_42 | Unknown | 2020-01-23 | Centers for Disease Control, R.O.C. (Taiwan) | Centers for Disease Control, R.O.C. (Taiwan) | Ji-Rong Yang, Yu-Chi Lin, Jung-Jung Mu, Ming-Tsan Liu, Shu-Ying Li |
| 91. | EPI_ISL_406030 | BetaCoV/Shenzhen/HKU-SZ-002/2020 | Asia/China/ Guangdong/ Shenzhen | Hap_15 | Unknown | 2020-01-10 | The University of Hong Kong - Shenzhen Hospital | Li Ka Shing Faculty of Medicine, The University of Hong Kong | Chan,J.F.-W., Yuan,S., Kok,K.H., To,K.K.-W., Chu,H., Yang,J., Xing,F., Liu,J., Yip,C.C.-Y., Poon,R.W.-S., Tsai,H.W., Lo,S.K.-F., Chan,K.H., Poon,V.K.-M., Chan,W.M., Ip,J.D., Cai,J.P., Cheng,V.C.-C., Chen,H., Hui,C.K.-M. and Yuen,K.Y. |
| 92. | EPI_ISL_405839 | BetaCoV/Shenzhen/HKU-SZ-005/2020 | Asia/China/ Guangdong/ Shenzhen | Hap_17 | Unknown | 2020-01-11 | The University of Hong Kong - Shenzhen Hospital | Li Ka Shing Faculty of Medicine, The University of Hong Kong | Chan,J.F.-W., Yuan,S., Kok,K.H., To,K.K.-W., Chu,H., Yang,J., Xing,F., Liu,J., Yip,C.C.-Y., Poon,R.W.-S., Tsai,H.W., Lo,S.K.-F., Chan,K.H., Poon,V.K.-M., Chan,W.M., Ip,J.D., Cai,J.P., Cheng,V.C.-C., Chen,H., Hui,C.K.-M. and Yuen,K.Y. |
| 93. | EPI_ISL_404895 | BetaCoV/USA/WA1/2020 | North America/ USA/ Washington/ Snohomish County | Hap_31 | Unknown | 2020-01-19 | Providence Regional Medical Center | Division of Viral Diseases, Centers for Disease Control and Prevention | Queen,K., Tao,Y., Li,Y., Paden,C.R., Lu,X., Zhang,J., Gerber,S.I., Lindstrom,S., Tong,S. |
| 94. | EPI_ISL_404253  ※ | BetaCoV/USA/IL1/2020 | North America/ USA/Illinois/ Chicago |  | Unknown | 2020-01-21 | IL Department of Public Health Chicago Laboratory | Pathogen Discovery, Respiratory Viruses Branch, Division of Viral Diseases, Centers for Dieases Control and Prevention | Ying Tao, Krista Queen, Clinton R. Paden, Jing Zhang, Yan Li, Anna Uehara, Xiaoyan Lu, Brian Lynch, Senthil Kumar K. Sakthivel, Brett L. Whitaker, Shifaq Kamili, Lijuan Wang, Janna' R. Murray, Susan I. Gerber, Stephen Lindstrom, Suxiang Tong |
| 95. | EPI_ISL_404228 | BetaCoV/Zhejiang/WZ-02/2020 | Asia/China/ Zhejiang | Hap_3 | Unknown | 2020-01-17 | Zhejiang Provincial Center for Disease Control and Prevention | Department of Microbiology, Zhejiang Provincial Center for Disease Control and Prevention | Yanjun Zhang, Yin Chen, Haiyan Mao, Junhang Pan, Xiuyu Lou, Yiyu Lu, Juying Yan, Hanping Zhu, Jian Gao, Yan Feng, Yi Sun, Hao Yan, Zhen Li, Yisheng Sun, Liming Gong, Qiong Ge, Wen Shi, Xinying Wang, Wenwu Yao, Zhangnv Yang, Fang Xu, Chen Chen, Enfu Chen, Zhen Wang, Zhiping Chen, Jianmin Jiang, Chonggao Hu |
| 96. | EPI_ISL_404227 | BetaCoV/Zhejiang/WZ-01/2020 | Asia/China/ Zhejiang | Hap_23 | Unknown | 2020-01-16 | Zhejiang Provincial Center for Disease Control and Prevention | Department of Microbiology, Zhejiang Provincial Center for Disease Control and Prevention | Yin Chen, Yanjun Zhang, Haiyan Mao, Junhang Pan, Xiuyu Lou, Yiyu Lu, Juying Yan, Hanping Zhu, Jian Gao, Yan Feng, Yi Sun, Hao Yan, Zhen Li, Yisheng Sun, Liming Gong, Qiong Ge, Wen Shi, Xinying Wang, Wenwu Yao, Zhangnv Yang, Fang Xu, Chen Chen, Enfu Chen, Zhen Wang, Zhiping Chen, Jianmin Jiang, Chonggao Hu |
| 97. | EPI_ISL_403963 | BetaCoV/Nonthaburi/74/2020 | Asia/Thailand/ Nonthaburi | Hap_3 | Unknown | 2020-01-13 | Bamrasnaradura Hospital | 1. Department of Medical Sciences, Ministry of Public Health, Thailand 2. Thai Red Cross Emerging Infectious Diseases - Health Science Centre 3. Department of Disease Control, Ministry of Public Health, Thailand | Pilailuk,Okada; Siripaporn,Phuygun; Thanutsapa,Thanadachakul; Supaporn,Wacharapluesadee; Sittiporn,Parnmen; Warawan,Wongboot; Sunthareeya,Waicharoen; Rome,Buathong; Malinee,Chittaganpitch; Nanthawan,Mekha |
| 98. | EPI_ISL_403962 | BetaCoV/Nonthaburi/61/2020 | Asia/Thailand/ Nonthaburi | Hap_3 | Unknown | 2020-01-08 | Bamrasnaradura Hospital | 1. Department of Medical Sciences, Ministry of Public Health, Thailand 2. Thai Red Cross Emerging Infectious Diseases - Health Science Centre 3. Department of Disease Control, Ministry of Public Health, Thailand | Pilailuk,Okada; Siripaporn,Phuygun; Thanutsapa,Thanadachakul; Supaporn,Wacharapluesadee; Sittiporn,Parnmen; Warawan,Wongboot; Sunthareeya,Waicharoen; Rome,Buathong; Malinee,Chittaganpitch; Nanthawan,Mekha |
| 99. | EPI_ISL_403937 | BetaCoV/Guangdong/20SF040/2020 | Asia/China/ Guangdong/ Zhuhai | Hap_26 | Unknown | 2020-01-18 | Guangdong Provincial Center for Diseases Control and Prevention; Guangdong Provincial Public Health | Department of Microbiology, Guangdong Provincial Center for Diseases Control and Prevention | Min Kang, Jie Wu, Jing Lu, Tao Liu, Baisheng Li, Shujiang Mei, Feng Ruan, Lifeng Lin, Changwen Ke, Haojie Zhong, Yingtao Zhang, Lirong Zou, Xuguang Chen, Qi Zhu, Jianpeng Xiao, Jianxiang Geng, Zhe Liu, Jianxiong Hu, Weilin Zeng, Xing Li, Yuhuang Liao, Xiujuan Tang, Songjian Xiao, Ying Wang, Yingchao Song, Xue Zhuang, Lijun Liang, Guanhao He, Huihong Deng, Tie Song, Jianfeng He, Wenjun Ma |
| 100. | EPI_ISL_403936 | BetaCoV/Guangdong/20SF028/2020 | Asia/China/ Guangdong/ Zhuhai | Hap_26 | Unknown | 2020-01-17 | Guangdong Provincial Center for Diseases Control and Prevention; Guangdong Provincial Public Health | Department of Microbiology, Guangdong Provincial Center for Diseases Control and Prevention | Min Kang, Jie Wu, Jing Lu, Tao Liu, Baisheng Li, Shujiang Mei, Feng Ruan, Lifeng Lin, Changwen Ke, Haojie Zhong, Yingtao Zhang, Lirong Zou, Xuguang Chen, Qi Zhu, Jianpeng Xiao, Jianxiang Geng, Zhe Liu, Jianxiong Hu, Weilin Zeng, Xing Li, Yuhuang Liao, Xiujuan Tang, Songjian Xiao, Ying Wang, Yingchao Song, Xue Zhuang, Lijun Liang, Guanhao He, Huihong Deng, Tie Song, Jianfeng He, Wenjun Ma |
| 101. | EPI_ISL_403935 | BetaCoV/Guangdong/20SF025/2020 | Asia/China/ Guangdong/ Shenzhen | Hap_15 | Unknown | 2020-01-15 | Guangdong Provincial Center for Diseases Control and Prevention; Guangdong Provincial Public Health | Department of Microbiology, Guangdong Provincial Center for Diseases Control and Prevention | Min Kang, Jie Wu, Jing Lu, Tao Liu, Baisheng Li, Shujiang Mei, Feng Ruan, Lifeng Lin, Changwen Ke, Haojie Zhong, Yingtao Zhang, Lirong Zou, Xuguang Chen, Qi Zhu, Jianpeng Xiao, Jianxiang Geng, Zhe Liu, Jianxiong Hu, Weilin Zeng, Xing Li, Yuhuang Liao, Xiujuan Tang, Songjian Xiao, Ying Wang, Yingchao Song, Xue Zhuang, Lijun Liang, Guanhao He, Huihong Deng, Tie Song, Jianfeng He, Wenjun Ma |
| 102. | EPI_ISL_403934 | BetaCoV/Guangdong/20SF014/2020 | Asia/China/ Guandong/ Shenzhen | Hap_21 | Unknown | 2020-01-15 | Guangdong Provincial Center for Diseases Control and Prevention; Guangdong Provincial Public Health | Department of Microbiology, Guangdong Provincial Center for Diseases Control and Prevention | Min Kang, Jie Wu, Jing Lu, Tao Liu, Baisheng Li, Shujiang Mei, Feng Ruan, Lifeng Lin, Changwen Ke, Haojie Zhong, Yingtao Zhang, Lirong Zou, Xuguang Chen, Qi Zhu, Jianpeng Xiao, Jianxiang Geng, Zhe Liu, Jianxiong Hu, Weilin Zeng, Xing Li, Yuhuang Liao, Xiujuan Tang, Songjian Xiao, Ying Wang, Yingchao Song, Xue Zhuang, Lijun Liang, Guanhao He, Huihong Deng, Tie Song, Jianfeng He, Wenjun Ma |
| 103. | EPI_ISL_403933 | BetaCoV/Guangdong/20SF013/2020 | Asia/China/ Guandong/ Shenzhen | Hap_15 | Unknown | 2020-01-15 | Guangdong Provincial Center for Diseases Control and Prevention; Guangdong Provincial Public Health | Department of Microbiology, Guangdong Provincial Center for Diseases Control and Prevention | Min Kang, Jie Wu, Jing Lu, Tao Liu, Baisheng Li, Shujiang Mei, Feng Ruan, Lifeng Lin, Changwen Ke, Haojie Zhong, Yingtao Zhang, Lirong Zou, Xuguang Chen, Qi Zhu, Jianpeng Xiao, Jianxiang Geng, Zhe Liu, Jianxiong Hu, Weilin Zeng, Xing Li, Yuhuang Liao, Xiujuan Tang, Songjian Xiao, Ying Wang, Yingchao Song, Xue Zhuang, Lijun Liang, Guanhao He, Huihong Deng, Tie Song, Jianfeng He, Wenjun Ma |
| 104. | EPI_ISL_403932 | BetaCoV/Guangdong/20SF012/2020 | Asia/China/ Guandong/ Shenzhen | Hap_15 | Unknown | 2020-01-14 | Guangdong Provincial Center for Diseases Control and Prevention; Guangdong Provincial Public Health | Department of Microbiology, Guangdong Provincial Center for Diseases Control and Prevention | Min Kang, Jie Wu, Jing Lu, Tao Liu, Baisheng Li, Shujiang Mei, Feng Ruan, Lifeng Lin, Changwen Ke, Haojie Zhong, Yingtao Zhang, Lirong Zou, Xuguang Chen, Qi Zhu, Jianpeng Xiao, Jianxiang Geng, Zhe Liu, Jianxiong Hu, Weilin Zeng, Xing Li, Yuhuang Liao, Xiujuan Tang, Songjian Xiao, Ying Wang, Yingchao Song, Xue Zhuang, Lijun Liang, Guanhao He, Huihong Deng, Tie Song, Jianfeng He, Wenjun Ma |
| 105. | EPI_ISL_403931 | BetaCoV/Wuhan/IPBCAMS-WH-02/2019 | Asia/China/ Hubei / Wuhan | Hap_3 | Unknown | 2019-12-30 | Institute of Pathogen Biology, Chinese Academy of Medical Sciences & Peking Union Medical College | Institute of Pathogen Biology, Chinese Academy of Medical Sciences & Peking Union Medical College | Lili Ren, Jianwei Wang, Qi Jin, Zichun Xiang, Zhiqiang Wu, Chao Wu, Yiwei Liu |
| 106. | EPI_ISL_403930 | BetaCoV/Wuhan/IPBCAMS-WH-03/2019 | Asia/China/ Hubei/ Wuhan | Hap_9 | Unknown | 2019-12-30 | Institute of Pathogen Biology, Chinese Academy of Medical Sciences & Peking Union Medical College | Institute of Pathogen Biology, Chinese Academy of Medical Sciences & Peking Union Medical College | Lili Ren, Jianwei Wang, Qi Jin, Zichun Xiang, Zhiqiang Wu, Chao Wu, Yiwei Liu |
| 107. | EPI_ISL_403929 | BetaCoV/Wuhan/IPBCAMS-WH-04/2019 | Asia/China/ Hubei / Wuhan | Hap_3 | Yes | 2019-12-30 | Institute of Pathogen Biology, Chinese Academy of Medical Sciences & Peking Union Medical College | Institute of Pathogen Biology, Chinese Academy of Medical Sciences & Peking Union Medical College | Lili Ren, Jianwei Wang, Qi Jin, Zichun Xiang, Zhiqiang Wu, Chao Wu, Yiwei Liu |
| 108. | EPI_ISL_403928 | BetaCoV/Wuhan/IPBCAMS-WH-05/2020 | Asia/China/ Hubei / Wuhan | Hap_12 | Unknown | 2020-01-01 | Institute of Pathogen Biology, Chinese Academy of Medical Sciences & Peking Union Medical College | Institute of Pathogen Biology, Chinese Academy of Medical Sciences & Peking Union Medical College | Lili Ren, Jianwei Wang, Qi Jin, Zichun Xiang, Zhiqiang Wu, Chao Wu, Yiwei Liu |
| 109. | EPI_ISL_402132 | BetaCoV/Wuhan/HBCDC-HB-01/2019 | Asia/China/Hubei/ Wuhan | Hap_8 | Yes | 2019-12-30 | Wuhan Jinyintan Hospital | Hubei Provincial Center for Disease Control and Prevention | Bin Fang, Xiang Li, Xiao Yu, Linlin Liu, Bo Yang, Faxian Zhan, Guojun Ye, Xixiang Huo, Junqiang Xu, Bo Yu, Kun Cai, Jing Li, Yongzhong Jiang. |
| 110. | EPI_ISL_402131 | BetaCoV/bat/Yunnan/RaTG13/2013 | Asia/China/Yunnan /Pu'er |  | Unknown | 2013-07-24 | Wuhan Institute of Virology, Chinese Academy of Sciences | Wuhan Institute of Virology, Chinese Academy of Sciences | Yan Zhu, Ping Yu, Bei Li, Ben Hu, Hao-Rui Si, Xing-Lou Yang, Peng Zhou, Zheng-Li Shi |
| 111. | EPI_ISL_402130 | BetaCoV/Wuhan/WIV07/2019 | Asia/China / Hubei /Wuhan | Hap_7 | Yes | 2019-12-30 | Wuhan Jinyintan Hospital | Wuhan Institute of Virology, Chinese Academy of Sciences | Peng Zhou, Xing-Lou Yang, Ding-Yu Zhang, Lei Zhang, Yan Zhu, Hao-Rui Si, Zhengli Shi |
| 112. | EPI_ISL_402129 | BetaCoV/Wuhan/WIV06/2019 | Asia/China/ Hubei / Wuhan | Hap_3 | Unknown | 2019-12-30 | Wuhan Jinyintan Hospital | Wuhan Institute of Virology, Chinese Academy of Sciences | Peng Zhou, Xing-Lou Yang, Ding-Yu Zhang, Lei Zhang, Yan Zhu, Hao-Rui Si, Zhengli Shi |
| 113. | EPI_ISL_402128 | BetaCoV/Wuhan/WIV05/2019 | Asia/China/ Hubei / Wuhan | Hap_6 | Yes | 2019-12-30 | Wuhan Jinyintan Hospital | Wuhan Institute of Virology, Chinese Academy of Sciences | Peng Zhou, Xing-Lou Yang, Ding-Yu Zhang, Lei Zhang, Yan Zhu, Hao-Rui Si, Zhengli Shi |
| 114. | EPI_ISL_402127 | BetaCoV/Wuhan/WIV02/2019 | Asia/China/ Hubei / Wuhan | Hap_5 | Yes | 2019-12-30 | Wuhan Jinyintan Hospital | Wuhan Institute of Virology, Chinese Academy of Sciences | Peng Zhou, Xing-Lou Yang, Ding-Yu Zhang, Lei Zhang, Yan Zhu, Hao-Rui Si, Zhengli Shi |
| 115. | EPI_ISL_402125 | BetaCoV/Wuhan-Hu-1/2019 | Asia/China | Hap_3 | Yes | 2019-12-31 | unknown | National Institute for Communicable Disease Control and Prevention (ICDC) Chinese Center for Disease Control and Prevention (China CDC) | Zhang,Y.-Z., Wu,F., Chen,Y.-M., Pei,Y.-Y., Xu,L., Wang,W., Zhao,S., Yu,B., Hu,Y., Tao,Z.-W., Song,Z.-G., Tian,J.-H., Zhang,Y.-L., Liu,Y., Zheng,J.-J., Dai,F.-H., Wang,Q.-M., She,J.-L. and Zhu,T.-Y. |
| 116. | EPI_ISL_402124 | BetaCoV/Wuhan/WIV04/2019 | Asia/China/ Hubei / Wuhan | Hap_3 | Yes | 2019-12-30 | Wuhan Jinyintan Hospital | Wuhan Institute of Virology, Chinese Academy of Sciences | Peng Zhou, Xing-Lou Yang, Ding-Yu Zhang, Lei Zhang, Yan Zhu, Hao-Rui Si, Zhengli Shi |
| 117. | EPI_ISL_402123 | BetaCoV/Wuhan/IPBCAMS-WH-01/2019 | Asia/China/ Hubei / Wuhan | Hap_1 | Unknown | 2019-12-24 | Institute of Pathogen Biology, Chinese Academy of Medical Sciences & Peking Union Medical College | Institute of Pathogen Biology, Chinese Academy of Medical Sciences & Peking Union Medical College | Lili Ren, Jianwei Wang, Qi Jin, Zichun Xiang, Zhiqiang Wu, Chao Wu, Yiwei Liu |
| 118. | EPI_ISL_402121 | BetaCoV/Wuhan/IVDC-HB-05/2019 | Asia/China/ Hubei / Wuhan | Hap_4 | Yes | 2019-12-30 | National Institute for Viral Disease Control and Prevention, China CDC | National Institute for Viral Disease Control and Prevention, China CDC | Wenjie Tan，Xuejun Ma，Xiang Zhao，Wenling Wang，Yongzhong Jiang，Roujian Lu，Ji Wang，Peihua Niu, Weimin Zhou, Faxian Zhan，Weifeng Shi，Baoying Huang，Jun Liu，Li Zhao，Yao Meng，Fei Ye，Na Zhu, Xiaozhou He，Peipei Liu, Yang Li，Jing Chen，Wenbo Xu，George F. Gao，Guizhen Wu |
| 119. | EPI_ISL_402120 | BetaCoV/Wuhan/IVDC-HB-04/2020 | Asia/China/Hubei/ Wuhan | Hap_11 | Yes | 2020-01-01 | National Institute for Viral Disease Control and Prevention, China CDC | National Institute for Viral Disease Control and Prevention, China CDC | Wenjie Tan，Xiang Zhao，Wenling Wang，Xuejun Ma，Yongzhong Jiang，Roujian Lu，Ji Wang，Weimin Zhou，Peihua Niu，Peipei Liu，Faxian Zhan，Weifeng Shi，Baoying Huang，Jun Liu，Li Zhao，Yao Meng，Xiaozhou He，Fei Ye，Na Zhu，Yang Li，Jing Chen，Wenbo Xu，George F. Gao，Guizhen Wu |
| 120. | EPI_ISL_402119 | BetaCoV/Wuhan/IVDC-HB-01/2019 | Asia/China/Hubei/ Wuhan | Hap_3 | Yes | 2019-12-30 | National Institute for Viral Disease Control and Prevention, China CDC | National Institute for Viral Disease Control and Prevention, China CDC | Wenjie Tan，Xiang Zhao，Wenling Wang，Xuejun Ma，Yongzhong Jiang，Roujian Lu, Ji Wang, Weimin Zhou，Peihua Niu，Peipei Liu，Faxian Zhan，Weifeng Shi，Baoying Huang，Jun Liu，Li Zhao，Yao Meng，Xiaozhou He，Fei Ye，Na Zhu，Yang Li，Jing Chen，Wenbo Xu，George F. Gao，Guizhen Wu |
| 121. | EPI_ISL_411060 | BetaCoV/Fujian/8/2020 | Asia/China/ Fujian | Hap_31 | Unknown | 2020-01-21 | Fujian Center for Disease Control and Prevention | Fujian Center for Disease Control and Prevention | Chen Wei, Zhang Yanhua, He Wenxiang, Weng Yuwei |
| 122. | EPI_ISL_411066 | BetaCoV/Fujian/13/2020 | Asia/China/ Fujian | Hap_35 | Unknown | 2020-01-22 | Fujian Center for Disease Control and Prevention | Fujian Center for Disease Control and Prevention | Chen Wei, Zhang Yanhua, He Wenxiang, Weng Yuwei |
| 123. | EPI_ISL_411218 | BetaCoV/France/IDF0571/2020 | Europe/France/ Ile-de-France/ Paris | Hap_61 | Unknown | 2020-02-02 | Department of Infectious and Tropical Diseases, Bichat Claude Bernard Hospital, Paris | Laboratoire Virpath, CIRI U111, UCBL1, INSERM, CNRS, ENS Lyon | Olivier Terrier, Aurélien Traversier, Julien Fouret, Yazdan Yazdanpanah, Xavier Lescure, Catherine Legras-Lachuer, Alexandre Gaymard, Bruno Lina, Manuel Rosa-Calatrava |
| 124. | EPI_ISL_411219 | BetaCoV/France/IDF0386-islP1/2020 | Europe/France/ Ile-de-France/ Paris | Hap_41 | Unknown | 2020-01-28 | Department of Infectious and Tropical Diseases, Bichat Claude Bernard Hospital, Paris | Laboratoire Virpath, CIRI U111, UCBL1, INSERM, CNRS, ENS Lyon | Olivier Terrier, Aurélien Traversier, Julien Fouret, Yazdan Yazdanpanah, Xavier Lescure, Alexandre Gaymard, Bruno Lina, Manuel Rosa-Calatrava |
| 125. | EPI_ISL_411220 | BetaCoV/France/IDF0386-islP3/2020 | Europe/France/ Ile-de-France/ Paris | Hap_41 | Unknown | 2020-01-28 | Department of Infectious and Tropical Diseases, Bichat Claude Bernard Hospital, Paris | Laboratoire Virpath, CIRI U111, UCBL1, INSERM, CNRS, ENS Lyon | Olivier Terrier, Aurélien Traversier, Julien Fouret, Yazdan Yazdanpanah, Xavier Lescure, Alexandre Gaymard, Bruno Lina, Manuel Rosa-Calatrava |
| 126. | EPI_ISL_411902 | BetaCoV/Cambodia/0012/2020 | Asia/Cambodia/ Sihanoukville | Hap_52 | Unknown | 2020-01-27 | Virology Unit, Institut Pasteur du Cambodge. | Virology Unit, Institut Pasteur du Cambodge (Sequencing done by: Jessica E Manning/Jennifer A Bohl at Malaria and Vector Research Research Laboratory, National Institute of Allergy and Infectious Diseases and Vida Ahyong from Chan-Zuckerberg Biohub) | Erik A Karlsson, Jennifer A Bohl, Vida Ahyong, Veasna Duong, Philippe Dussart, Jessica E Manning. |
| 127. | EPI_ISL_411915 | BetaCoV/Taiwan/CGMH-CGU-01/2020 | Asia/Taiwan/  Taoyuan | Hap_3 | Unknown | 2020-01-25 | Laboratory Medicine | Department of Laboratory Medicine, Lin-Kou Chang Gung Memorial Hospital, Taoyuan, Taiwan. | Kuo-Chien Tsao, Yu-Nong Gong, Shu-Li Yang, Yi-Chun Li, Chung-Guei Huang, Yhu-Chering Huang, Shin-Ru Shih |
| 128. | EPI_ISL_411926 | BetaCoV/Taiwan/3/2020 | Asia/Taiwan/ Taipei | Hap_14 | Unknown | 2020-01-24 | Taiwan Centers for Disease Control | Taiwan Centers for Disease Control | Ji-Rong Yang, Yu-Chi-Lin, Jung-Jung Mu, Ming-Tsan-Liu |
| 129. | EPI_ISL_411927 | BetaCoV/Taiwan/4/2020 | Asia/Taiwan / Taipei | Hap_3 | Unknown | 2020-01-28 | Taiwan Centers for Disease Control | Taiwan Centers for Disease Control | Ji-Rong Yang, Yu-Chi-Lin, Jung-Jung Mu, Ming-Tsan-Liu |
| 130. | EPI_ISL_411929 | BetaCoV/South Korea/SNU01/2020 | Asia/South Korea | Hap_3 | Unknown | 2020-01-20 | unknown | Department of Clinical Diagnostics | Park,W.B., Kwon,N.-J., Choi,S.-J., Kang,C.K., Choe,P.G., Kim,J.Y., Yun,J., Lee,G.-W., Seong,M.-W., Kim,N., Seo,J.-S. and Oh,M.-D. |
| 131. | EPI_ISL_411950 | BetaCoV/Jiangsu/JS01/2020 | Asia/China/ Jiangsu | Hap_40 | Unknown | 2020-01-23 | NHC Key laboratory of Enteric Pathogenic Microbiology, Institute of Pathogenic Microbiology | Jiangsu Provincial Center for Disease Control & Prevention | Lunbiao Cui,Kangchen Zhao,Xiaojuan Zhu,Yiyue Ge,Tao Wu,Bin Wu,Yin Chen,Fengcai Zhu,Baoli Zhu,Ming Wu |
| 132. | EPI_ISL_411951 | BetaCoV/Sweden/01/2020 | Europe/Sweden | Hap_77 | Unknown | 2020-02-07 | unknown | Unit for Laboratory Development and Technology Transfer, Public Health Agency of Sweden | Bengner,M., Palmerus,M., Lindsjo,O., Lind Karlberg,M., Monteil,V., Appelberg,S., Brave,A., Muradrasoli,S. and Tegmark-Wisell,K. |
| 133. | EPI_ISL_411952 | BetaCoV/Jiangsu/JS02/2020 | Asia/China/ Jiangsu | Hap_45 | Unknown | 2020-01-24 | NHC Key laboratory of Enteric Pathogenic Microbiology, Institute of Pathogenic Microbiology | Jiangsu Provincial Center for Disease Control & Prevention | Kangchen Zhao, Xiaojuan Zhu, Lunbiao Cui, Tao Wu, Yiyue Ge, Bin Wu, Yin Chen, Fengcai Zhu, Baoli Zhu, Ming Wu |
| 134. | EPI_ISL_411953 | BetaCoV/Jiangsu/JS03/2020 | Asia/China/ Jiangsu | Hap_3 | Unknown | 2020-01-24 | NHC Key laboratory of Enteric Pathogenic Microbiology, Institute of Pathogenic Microbiology | Jiangsu Provincial Center for Disease Control & Prevention | Kangchen Zhao, Xiaojuan Zhu, Lunbiao Cui, Tao Wu, Yiyue Ge, Bin Wu, Yin Chen, Fengcai Zhu, Baoli Zhu, Ming Wu |
| 135. | EPI_ISL_411954 | BetaCoV/USA/CA7/2020 | North America/ USA /California | Hap_54 | Unknown | 2020-02-06 | California Department of Public Health | Pathogen Discovery, Respiratory Viruses Branch, Division of Viral Diseases, Centers for Dieases Control and Prevention | Krista Queen, Anna Uehara, Jing Zhang, Yan Li, Ying Tao, Clinton R. Paden, Haibin Wang, Shifaq Kamili, Xiaoyan Lu, Brian Lynch, Senthil Kumar K. Sakthivel, Brett L. Whitaker, Lijuan Wang, Janna' R. Murray, Susan I. Gerber, Stephen Lindstrom, Suxiang Tong |
| 136. | EPI_ISL_411955 | BetaCoV/USA/CA8/2020 | North America/ USA /California | Hap_10 | Unknown | 2020-02-10 | California Department of Public Health | Pathogen Discovery, Respiratory Viruses Branch, Division of Viral Diseases, Centers for Dieases Control and Prevention | Krista Queen, Anna Uehara, Jing Zhang, Yan Li, Ying Tao, Clinton R. Paden, Haibin Wang, Shifaq Kamili, Xiaoyan Lu, Brian Lynch, Senthil Kumar K. Sakthivel, Brett L. Whitaker, Lijuan Wang, Janna' R. Murray, Susan I. Gerber, Stephen Lindstrom, Suxiang Tong |
| 137. | EPI_ISL_411956 | BetaCoV/USA/TX1/2020 | North America/ USA/Texas | Hap_79 | Unknown | 2020-02-11 | Texas Department of State Health Services | Pathogen Discovery, Respiratory Viruses Branch, Division of Viral Diseases, Centers for Dieases Control and Prevention | Krista Queen, Anna Uehara, Jing Zhang, Yan Li, Ying Tao, Clinton R. Paden, Haibin Wang, Shifaq Kamili, Xiaoyan Lu, Brian Lynch, Senthil Kumar K. Sakthivel, Brett L. Whitaker, Lijuan Wang, Janna' R. Murray, Susan I. Gerber, Stephen Lindstrom, Suxiang Tong |
| 138. | EPI_ISL_411957 | BetaCoV/China/WH-09/2020 | Asia/China | Hap_3 | Unknown | 2020-01-08 | unknown | Key Laboratory of Human Diseases, Comparative Medicine, Institute of Laboratory Animal Science | Linlin,B., Lili,R., Shuran,G., Jiangning,L., Feifei,Q., Qi,L., Fengdi,L., Jing,X., Wei,D., Pin,Y., Yanfeng,X., Yajin,Q., Hong,G., Qiang,W., Mingya,L., Guanpeng,W., Shunyi,W., Zhiqi,S., Li,G., Lan,C., Conghui,W., Ying,W., Xinming,W., Yan,X., Qi,J. and Chuan,Q. |
| 139. | EPI_ISL_412978 | BetaCoV/Wuhan/HBCDC-HB-02/2020 | Asia/China/ Hubei / Wuhan | Hap_27 | No | 2020-01-08 | The Central Hospital Of Wuhan | Hubei Provincial Center for Disease Control and Prevention | Bin Fang, Xiang Li, Xiao Yu, Linlin Liu, Bo Yang, Faxian Zhan, Guojun Ye, Xixiang Huo, Junqiang Xu, Bo Yu, Kun Cai, Jing Li, Yongzhong Jiang. |
| 140. | EPI_ISL_412979 | BetaCoV/Wuhan/HBCDC-HB-03/2020 | Asia/China/ Hubei / Wuhan | Hap_14 | No^δ^ | 2020-01-18 | Union Hospital of Tongji Medical College, Huazhong University of Science and Technology | Hubei Provincial Center for Disease Control and Prevention | Bin Fang, Xiang Li, Xiao Yu, Linlin Liu, Bo Yang, Faxian Zhan, Guojun Ye, Xixiang Huo, Junqiang Xu, Bo Yu, Kun Cai, Jing Li, Yongzhong Jiang. |
| 141. | EPI_ISL_412980 | BetaCoV/Wuhan/HBCDC-HB-04/2020 | Asia/China/ Hubei / Wuhan | Hap_28 | No | 2020-01-18 | Union Hospital of Tongji Medical College, Huazhong University of Science and Technology | Hubei Provincial Center for Disease Control and Prevention | Bin Fang, Xiang Li, Xiao Yu, Linlin Liu, Bo Yang, Faxian Zhan, Guojun Ye, Xixiang Huo, Junqiang Xu, Bo Yu, Kun Cai, Jing Li, Yongzhong Jiang. |
| 142. | EPI_ISL_412898 | BetaCoV/Wuhan/HBCDC-HB-02/2019 | Asia/China/ Hubei/ Wuhan | Hap_10 | Yes | 2019-12-30 | Wuhan Jinyintan Hospital | Hubei Provincial Center for Disease Control and Prevention | Bin Fang, Xiang Li, Xiao Yu, Linlin Liu, Bo Yang, Faxian Zhan, Guojun Ye, Xixiang Huo, Junqiang Xu, Bo Yu, Kun Cai, Jing Li, Yongzhong Jiang. |
| 143. | EPI_ISL_412899 | BetaCoV/Wuhan/HBCDC-HB-03/2019 | Asia/China/ Hubei / Wuhan | Hap_3 | Yes | 2019-12-30 | Wuhan Jinyintan Hospital | Hubei Provincial Center for Disease Control and Prevention | Bin Fang, Xiang Li, Xiao Yu, Linlin Liu, Bo Yang, Faxian Zhan, Guojun Ye, Xixiang Huo, Junqiang Xu, Bo Yu, Kun Cai, Jing Li, Yongzhong Jiang. |
| 144. | EPI_ISL_412981 | BetaCoV/Wuhan/HBCDC-HB-05/2020 | Asia/China/ Hubei / Wuhan | Hap_29 | No | 2020-01-18 | CR&WISCO GENERAL HOSPITAL | Hubei Provincial Center for Disease Control and Prevention | Bin Fang, Xiang Li, Xiao Yu, Linlin Liu, Bo Yang, Faxian Zhan, Guojun Ye, Xixiang Huo, Junqiang Xu, Bo Yu, Kun Cai, Jing Li, Yongzhong Jiang. |
| 145. | EPI_ISL_412983 | BetaCoV/Tianmen/HBCDC-HB-07/2020 | Asia/China/Hubei/ Tianmen | Hap_78 | No | 2020-02-08 | Tianmen Center for Disease Control and Prevention | Hubei Provincial Center for Disease Control and Prevention | Bin Fang, Xiang Li, Xiao Yu, Linlin Liu, Bo Yang, Faxian Zhan, Guojun Ye, Xixiang Huo, Junqiang Xu, Bo Yu, Kun Cai, Jing Li, YiFa Zhu, Yangyang Tao,Xierong Li,Yongzhong Jiang. |
| 146. | EPI_ISL_412459 | BetaCoV/Jingzhou/HBCDC-HB-01/2020 | Asia/China/Hubei/ Jingzhou | Hap_16 | Yes | 2020-01-08 | Jingzhou Center for Disease Control and Prevention | Hubei Provincial Center for Disease Control and Prevention | Bin Fang, Xiang Li, Xiao Yu, Linlin Liu, Bo Yang, Faxian Zhan, Guojun Ye, Xixiang Huo, Junqiang Xu, Bo Yu, Kun Cai, Jing Li, Maoyi Chen,Jie Hu, Chunlin Mao, Yongzhong Jiang. |
| 147. | EPI_ISL_412982 | BetaCoV/Wuhan/HBCDC-HB-06/2020 | Asia/China/ Hubei / Wuhan | Hap_76 | No | 2020-02-07 | Wuhan Lung Hospital | Hubei Provincial Center for Disease Control and Prevention | Bin Fang, Xiang Li, Xiao Yu, Linlin Liu, Bo Yang, Faxian Zhan, Guojun Ye, Xixiang Huo, Junqiang Xu, Bo Yu, Kun Cai, Jing Li, Yongzhong Jiang. |

Note: δ, the host live in a residential area about 2 kilometers from the Huanan seafood market; ※，sequences that not used in network analysis.
