## Supplemental Table 3 for "Genome-wide data inferring the evolution and population demography of the novel pneumonia coronavirus (SARS-CoV-2)"

| Start date | End data | Sequence Number | Haplotype number | π | Hd | Tajima's D | Fu's FS |
| --- | --- | --- | --- | --- | --- | --- | --- |
| 24^th^ 12, 2019 | 30^th^ 12, 2019 | 15 | 10 | 0.00007 | 0.857 | -2.16252 | -5.85468 |
| 24^th^ 12, 2019 | 6^th^ 1, 2020 | 24 | 14 | 0.00007 | 0.801 | -2.48655 | -10.46702 |
| 24^th^ 12, 2019 | 13^th^ 1, 2020 | 34 | 20 | 0.00013 | 0.836 | -2.64195 | -10.66549 |
| 24^th^ 12, 2019 | 20^th^ 1, 2020 | 59 | 36 | 0.00017 | 0.904 | -2.72675 | -25.37821 |
| 24^th^ 12, 2019 | 27^th^ 1, 2020 | 96 | 55 | 0.00015 | 0.926 | -2.78097 | -25.72435 |
| 24^th^ 12, 2019 | 3^th^ 2, 2020 | 127 | 73 | 0.00015 | 0.945 | -2.76899 | -25.58251 |
| 24^th^ 12, 2019 | 11^th^ 2, 2020 | 140 | 80 | 0.00016 | 0.954 | -2.78621 | -25.46648 |

Table S3. Descriptive statistics of genomes of SARS-Cov-2.
