## Supplementary figures and images for "Genome-wide data inferring the evolution and population demography of the novel pneumonia coronavirus (SARS-CoV-2)"

### Supplemental Figure 1

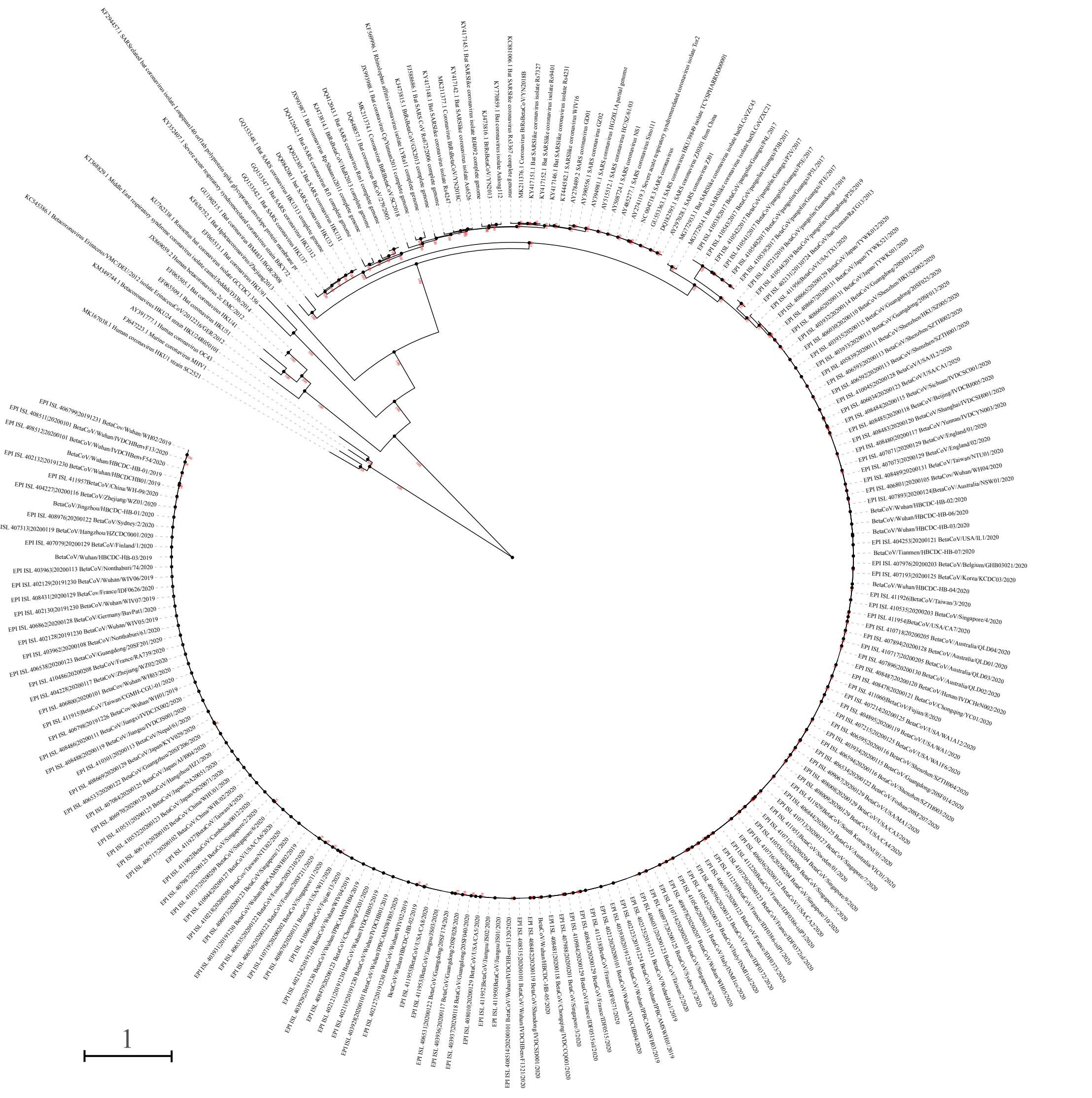
